# Arabidopsis AAR2, a conserved splicing factor in eukaryotes, acts in microRNA biogenesis

**DOI:** 10.1101/2022.06.26.497656

**Authors:** Lusheng Fan, Bin Gao, Ye Xu, Nora Flynn, Brandon Le, Chenjiang You, Shaofang Li, Natalia Achkar, Pablo A. Manavella, Zhenbiao Yang, Xuemei Chen

**Affiliations:** FAFU-UCR Joint Center for Horticultural Biology and Metabolomics Center, Haixia Institute of Science and Technology, Fujian Agriculture and Forestry University, Fuzhou, Fujian, China; Department of Botany and Plant Sciences, Institute of Integrative Genome Biology, University of California, Riverside, CA, USA; State Key Laboratory of Genetic Engineering and Collaborative Innovation Center of Genetics and Development, Institute of Plant Biology, School of Life Sciences, Fudan University, Shanghai 200438; State Key Laboratory for Biology of Plant Diseases and Insect Pests, Institute of Plant Protection, Chinese Academy of Agricultural Sciences, Beijing 100193, China; Instituto de Agrobiotecnologίa del Litoral (CONICET-UNL-FBCB), Santa Fe 3000, Argentina

## Abstract

MicroRNAs (miRNAs) play an essential role in plant growth and development, and as such, their biogenesis is fine-tuned via regulation of the core microprocessor components. Here, we report that *Arabidopsis* AAR2, a homolog of a U5 snRNP assembly factor in yeast and humans, not only acts in splicing but also promotes miRNA biogenesis. AAR2 interacts with the microprocessor component HYPONASTIC LEAVES1 (HYL1) in the cytoplasm, nucleus and dicing bodies. In *aar2* mutants, abundance of nonphosphorylated HYL1, the active form of HYL1, and the number of HYL1-labeled dicing bodies are reduced. Primary miRNA (pri-miRNA) accumulation is compromised despite normal promoter activities of *MIR* genes in *aar2* mutants. RNA decay assays show that the *aar2-1* mutation leads to faster degradation of pri-miRNAs in a HYL1-dependent manner, which reveals a previously unknown and negative role of HYL1 in miRNA biogenesis. Taken together, our findings reveal a dual role of AAR2 in miRNA biogenesis and pre-mRNA splicing.

**Significance:** In yeast and humans, AAR2 is involved in pre-mRNA splicing through regulating U5 snRNP assembly. This study shows that *Arabidopsis* AAR2 promotes microRNA (miRNA) accumulation in addition to its conserved role in pre-mRNA splicing. AAR2 is associated with the microprocessor component HYL1 and promotes its dephosphorylation to produce the active form in miRNA biogenesis. The study also reveals a previously unknown role of HYL1 in causing the degradation of the primary precursors to miRNAs (pri-miRNAs) and a role of AAR2 in protecting pri-miRNAs from HYL1-depedent degradation. Taken together, our findings provide new insights into the role of a conserved splicing factor in miRNA biogenesis in plants.

## Introduction

Small RNAs of 21-24 nucleotides (nt), including microRNAs (miRNAs) and small interfering RNAs (siRNAs), are involved in a wide range of biological processes (1). miRNAs are derived from primary transcripts with partially complementary fold-backs while siRNAs are generated from long, perfect or nearly perfect double-stranded RNAs. Both siRNAs and miRNAs are bound by Argonaute proteins to form RNA-induced silencing complexes (RISCs), which regulate gene expression either through transcriptional gene silencing or post-transcriptional gene silencing in a sequence-specific manner (2, 3).

In plants, most miRNA genes (*MIR*) are transcribed as independent transcription units to produce primary miRNAs (pri-miRNAs) that contain a stem loop structure (4). In *Arabidopsis*, a pri-miRNA is processed in the nucleus by DICER-LIKE1 (DCL1) into a miRNA/miRNA* duplex, which is 2’-*O*-methlyated at the 3’ terminal nucleotides by HUA ENHACER1 (HEN1); the duplex is bound by AGO1 followed by RISC formation upon the ejection of the miRNA* strand; miRISCs (miRNA-RISC complexes) are then exported to the cytoplasm and repress the expression of target genes via transcript cleavage or translational repression (2, 5-7). In Arabidopsis, several miRNAs target noncoding transcripts from *TRANS-ACTING SIRNA* (*TAS*) loci for cleavage and trigger the production of siRNAs from the cleavage fragments (1). These siRNAs are termed ta-siRNAs as they regulate target genes *in trans* (8).

The precise and efficient processing of pri-miRNAs by DCL1 is enhanced by the double-stranded RNA-binding protein HYPONASTIC LEAVES 1 (HYL1) and the zinc finger protein SERRATE (SE); the three proteins form a protein complex called the microprocessor (9, 10). In *Arabidopsis, dcl1* and *se* null mutations cause embryonic lethality, and *hyl1* null alleles lead to strong developmental defects (11-13). Consistent with their coordinated function in pri-miRNA processing, these three proteins co-localize in nuclear membrane-less speckles, known as Dicing bodies (D-bodies), as well as in the nucleoplasm, both of which are thought to be sites of pri-miRNA processing (14, 15). SE contains three intrinsically disordered regions that mediate phase separation, which plays a role in D-body formation (16). Besides its role in miRNA biogenesis, SE is also involved in pre-mRNA splicing. Like SE (17), other factors first found to play a role in splicing, such as CAP-BINDING PROTEIN 80 (CBP80), CAP-BINDING PROTEIN 20 (CBP20), STABLILIZED1 (STA1), proteins in the MOS4-associated complex (MAC) including MAC3A, MAC3B, PRL1, PRL2, and Serrate-Associated Protein 1 (SEAP1), were shown to be required for miRNA biogenesis (18-21). Conversely, several factors first found to play a role in miRNA biogenesis, such as Protein Phosphatase 4 (PP4) Regulatory Subunit 3 (22), THP1 in the TREX-2 complex (23), and RBV (24), were also found to act in the pre-mRNA splicing for subsets of genes.

Cofactors of the RNAseIII enzymes involved miRNA biogenesis are regulated to fine-tune miRNA production (25). In animals, TRBP and DGCR8, cofactors of Dicer and Drosha, respectively, are phosphorylated by mitogen-activated protein kinase (MAPK), which stabilizes the respective protein complexes to promote miRNA biogenesis (26, 27). However, in *Arabidopsis*, de-phosphorylation of HYL1 by CPL1 and CPL2 facilitates the localization of HYL1 in D-bodies and enhances miRNA biogenesis (28). MPK3 was identified as a kinase phosphorylating HYL1 in both rice and *Arabidopsis* and a mutation in Arabidopsis *MPK3* led to increased miRNA accumulation (29). SMEK1 (Suppressor of MEK 1) inhibits MAPK-mediated HYL1 phosphorylation and promotes PP4-mediated HYL1 de-phosphorylation, which is required for miRNA biogenesis (30). The core photomorphogenic regulator COP1 modulates the degradation of HYL1 in response to the light-to-dark transition (31). The phosphorylated form of HYL1 is inactive and less susceptible to degradation induced by extended periods of darkness. Upon light restoration, HYL1 is quickly reactivated by dephosphorylation, which in turn enables miRNA production and the proper expression of developmentally important genes (32). In response to ABA treatment, SnRK2 kinases phosphorylate HYL1, which is important for the stability of HYL1 and miRNA accumulation (33). These findings suggest that dephosphorylated HYL1 is the active form of the protein in miRNA biogenesis, but phosphorylated HYL1 is more stable and may serve as a reservoir from which dephosphorylated HYL1 can be produced.

SE is another important protein that assists DCL1 in pri-miRNA processing. Intriguingly, while SE is considered a component of the microprocessor, recent studies also point to a negative effect of SE in miRNA biogenesis. CHR2, an ATPase subunit of the SWI/SNF chromatin-remodeling complex, accesses pri-miRNAs through its interaction with SE and remodels the structures of pri-miRNAs to inhibit pri-miRNA processing (34). A MAC component, MAC5A, binds the stem-loop region of pri-miRNAs and protects pri-miRNAs from degradation, thus promoting miRNA biogenesis; the degradation of pri-miRNAs in *mac5a* mutants is dependent on SE and partially caused by 5’-to-3’ exoribonucleases XRN2 and XRN3 (35). How the positive and negative roles of SE in miRNA biogenesis are balanced is as yet unknown. SE is phosphorylated by pre-mRNA processing 4 kinase A (PRP4KA), leading to lower binding affinity to HYL1 as well as higher susceptibility to degradation by the 20S proteasome. Phosphorylation of SE via PRP4KA may define a regulatory mechanism to clear excess SE to maintain its proper levels and promote miRNA production (36). Whether the other microprocessor component, HYL1, has a negative role in miRNA biogenesis is unknown.

In this study, we performed a genetic screen in *Arabidopsis* using *SUC2:atasi-SUL* as a reporter to identify genes involved in ta-siRNA or miRNA biogenesis and activity (37). In the *SUC2:atasi-SUL* transgenic line, atasi-SUL, an artificial ta-siRNA, is engineered in the backbone of *TAS1* such that its biogenesis is triggered by miR173 as for endogenous ta-siRNAs from this locus (37). atasi-SUL represses the expression of the *SULFUR* (*SUL*) gene in mesophyll cells leading to leaf bleaching. Through this mutagenesis screen, we isolated a mutant showing a global reduction in both ta-siRNA and miRNA production. This mutation is in an uncharacterized gene encoding a protein homologous to AAR2, a U5 small nuclear ribonucleoprotein (snRNP) particle assembly factor in yeast and humans (38, 39). We show that *aar2* mutations reduce pri-miRNA levels without affecting the transcription of the corresponding *MIR* genes. The reduction of pri-miRNA levels in *aar2* is caused by pri-miRNA degradation that is dependent on HYL1, but not SE, indicating that HYL1 also has a negative effect in miRNA biogenesis. AAR2 associates with HYL1 and promotes its localization to D-bodies. The ratio between non-phosphorylated and phosphorylated HYL1 is decreased in the *aar2* mutant, suggesting that AAR2 promotes HYL1 dephosphorylation. Finally, AAR2 is also involved in pre-mRNA splicing for a subset of introns. Our findings suggest that in addition to its conserved role in pre-mRNA splicing, AAR2 is involved in miRNA biogenesis by preventing HYL1-dependent pri-miRNA degradation and promoting HYL1 dephosphorylation.

## RESULTS

### Isolation of a mutant with pleiotropic developmental defects and a global reduction in miRNA accumulation

To identify genes required for the biogenesis and/or activity of ta-siRNAs, we took advantage of a visual reporter of ta-siRNA activity, *SUC2::atasi-SUL* (*atasi-SUL*) (37), to perform an ethyl methanesulfonate mutagenesis screen. In the *atasi-SUL* transgenic line, an artificial ta-siRNA targeting the chlorophyll biosynthetic gene *SULFUR* (*SUL*, also known as *CHLORINA42*) leads to the yellowing of leaves. We isolated a suppressor mutant with green leaves, a phenotype indicative of compromised atasi-SUL biogenesis or activity (Fig. 1A). To identify the mutation, we backcrossed the mutant with the parental line *atasi-SUL* and conducted whole-genome re-sequencing using pooled F2 segregants with the mutant phenotype. A G^2400^-to-A mutation in AT1G66510 was identified, which introduced a premature stop codon in the open reading frame (*SI Appendix* Fig. S1A). This gene (referred to as *AAR2* hereafter) encodes a homolog to AAR2 that is involved in U5 snRNP assembly and consequently pre-mRNA splicing in yeast and humans (38). To examine the developmental phenotypes caused by the *aar2* mutation, we crossed this mutant with wild-type plants and obtained the *aar2-1* allele without the *atasi-SUL* transgene. We also acquired an *aar2* T-DNA insertion allele, SALK117746 (designated as *aar2-2* hereafter) (*SI Appendix* Fig. S1A). RNA sequencing (RNA-seq) with these two mutants revealed reduced levels of *AAR2* transcripts as compared with wild type (*SI Appendix* Fig. S1B). *aar2-2* plants showed similar developmental phenotypes to *aar2-1* plants, including reduced plant size, shorter primary roots, lower number of lateral roots and fewer stamens (Fig. 1A and *SI Appendix* Fig. S2 A-J). Both *aar2* mutants had shorter hypocotyls as compared to wild type when grown in darkness, and a similar phenotype was also observed for *hyl1-2* (*SI Appendix* Fig. S2 K-L). To determine whether the *aar2-1* mutation was responsible for the reduced leaf yellowing in *aar2-1 atasi-SUL*, we generated a construct in which a genomic fragment containing the promoter and coding region of *AAR2* was fused with yellow fluorescent protein (YFP) and transformed the construct into *aar2-1 atasi-SUL*. The transgene fully rescued the leaf color phenotype and the developmental defects of *aar2-1 atasi-SUL* (Fig. 1A). These results demonstrated that the *aar2-1* mutation is responsible for these phenotypes.

**Fig. 1.**
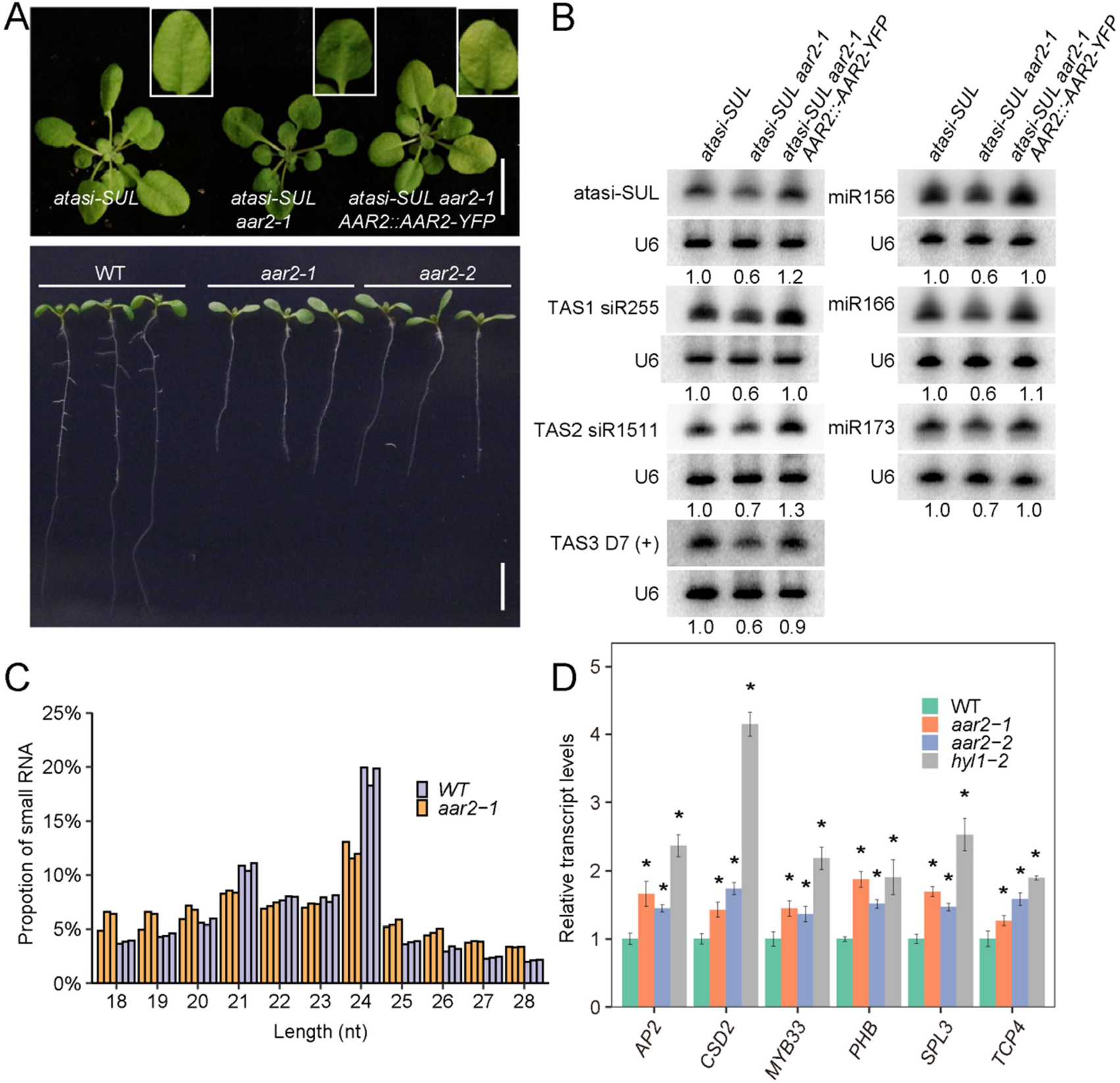
Isolation of a mutant with pleiotropic developmental defects and a global reduction in miRNA accumulation. (*A*) Phenotypes of *atasi-SUL, atasi-SUL aar2-1, AAR2::AAR2-YFP atasi-SUL aar2-1*, wild type (WT), *aar2-1* and *aar2-2* plants. Images of rosettes and roots were taken from 3-week-old (upper panel) and 9-day-old (lower panel) plants. Scale bar, 10 mm (upper panel) and 5 mm (lower panel). (*B*) RNA gel blot analysis showing reduced accumulation of *atasi-SUL*, endogenous ta-siRNAs and miRNAs in the *atasi-SUL aar2-1* mutant. Note that expression of the *AAR2::AAR2-YFP* transgene fully rescued the ta-siRNA and miRNA accumulation defects. U6 was used as the loading control. The numbers below the blots represent the relative amounts of ta-siRNAs and miRNAs. (*C*) The length distribution of mapped small RNA reads from small RNA-seq with WT and *aar2-1*. (*D*) Determination of miRNA target transcript levels in WT, *aar2-1, aar2-2* and *hyl1-2* by qRT-PCR. Transcript levels were normalized to *UBQ5* and then calculated relative to those in WT. Error bars represent SD calculated from three independent replicates. Asterisks indicate significant difference (Student’s t-test, *P < 0.05).

Given that the *aar2-1 atasi-SUL* mutant showed compromised atasi-SUL activity, we performed RNA gel blot analysis to determine the levels of atasi-SUL as well as endogenous ta-siRNAs in the *aar2-1 atasi-SUL* mutant. Both atasi-SUL and endogenous ta-siRNAs showed reduced accumulation in *aar2-1 atasi-SUL* as compared to *atasi-SUL* (Fig 1B). As ta-siRNA production is dependent on their trigger miRNAs, we further examined the levels of miR173 and miR390, the trigger miRNAs for ta-siRNA production from *TAS1, TAS2* and *TAS3*, as well as the levels of other endogenous miRNAs. All examined miRNAs showed moderate reduction in abundance in *aar2-1 atasi-SUL* (Fig 1B and *SI Appendix* Fig. S3A). These results indicate that the *aar2-1* mutation impaired miRNA production and indirectly affected ta-siRNA biogenesis. Moreover, the *AAR2-YFP* transgene fully rescued the ta-siRNA and miRNA production defects in *aar2-1*, further confirming that AAR2 is required for ta-siRNA and miRNA biogenesis (Fig. 1B). To assess the global profiles of small RNAs in the *aar2* mutant, we performed small RNA-seq with *aar2-1* and wild-type plants. Three biological replicates for each genotype were included and they were highly correlated (*SI Appendix* Fig. S3B). A significant global reduction in 21-nt to 24-nt small RNAs, which consist of most endogenous miRNAs and siRNAs, was found in *aar2-1* (Fig. 1C and *SI Appendix* Fig. S3C). miRNAs also showed a global reduction in *aar2-1* with 39 miRNAs showing a significant decrease in abundance (*SI Appendix* Fig. S3D and Dataset S1). In plants, pri-miRNAs are processed in either a base-to-loop or a loop-to-base manner (40, 41). We found that the reduction in miRNA abundance in *aar2-1* was regardless of the direction of pri-miRNA processing (*SI Appendix* Fig. S4A). When compared to all detected miRNAs, the 39 down-regulated miRNAs in *aar2-1* did not show a significant difference in the lengths of pri-miRNAs (*SI Appendix* Fig. S4 C). In comparison to 39 randomly selected miRNAs, the 39 down-regulated miRNAs in *aar2-1* represent significantly fewer miRNA families (*SI Appendix* Fig. S4 C), suggesting that some affected miRNAs belong to evolutionarily old miRNA families with more than one member (42). Consistent with such “old” miRNAs having higher abundance (42), the 39 miRNAs show higher levels as compared to all detected miRNAs in wild type (*SI Appendix* Fig. S4 D). The reduction in the levels of miRNAs in *aar2* mutants correlated with an increase in known miRNA-targeted mRNAs, as expected for a miRNA deficient mutant (Fig. 1D). To test the possibility that AAR2 affects miRNA production through promoting the expression of key genes in miRNA biogenesis, we examined the transcript or protein levels of known players in miRNA biogenesis or activity, and no significant changes were found in the *aar2* mutants (*SI Appendix* Fig. S5).

### AAR2 associates with microprocessor components and is required for the localization of HYL1 in D-bodies

To further investigate the molecular functions of the AAR2 protein, we first examined its subcellular localization. In the *AAR2::AAR2-YFP* transgenic line, AAR2-YFP signals were ubiquitous in root and leaf cells and were present in both the nucleus and the cytoplasm (*SI Appendix* Fig. S6). In root tip cells of the *HYL1-YFP AAR2-mCherry* transgenic line, AAR2-mCherry co-localized with HYL1-YFP in the nucleoplasm but was absent from the HYL1-YFP-labeled D-bodies (Fig. 2A). However, we found that HYL1-YFP-labeled D-bodies were significantly reduced in number in *aar2-1* as compared to wild type (Fig. 2B-D), suggesting that AAR2 promotes the localization of HYL1 in D-bodies. Given the fact that DCL1, HYL1 and SE colocalize in D-bodies (14) and *aar2-1* is compromised in the D-body localization of HYL1, we speculated that AAR2 may be associated with the microprocessor and act as an accessory component. Thus, we performed bimolecular fluorescence complementation (BiFC) analysis to test the association of AAR2 with DCL1, HYL1 and SE. In this assay, we fused AAR2 with an N-terminal fragment of YFP at its C-terminus (AAR2-nYFP) and co-expressed it with DCL1, HYL1, and SE tagged with a C-terminal fragment of YFP at their C-termini in *Nicotiana benthamiana*. We also included HYL1-nYFP and DCL1-cYFP as a positive control, which have been shown to associate with each other in the nucleus (14). As expected, co-expression of HYL1-nYFP and DCL1-cYFP generated YFP signals mainly in D-bodies in the nucleus (*SI Appendix* Fig. S7A). Similarly, co-expression of AAR2-nYFP with DCL1 and SE also generated YFP signals, but mainly in the nucleoplasm (*SI Appendix* Fig. S7A). Interestingly, co-expression of AAR2-nYFP with HYL1-cYFP gave rise to YFP signals in the cytoplasm, the nucleoplasm and D-bodies (*SI Appendix* Fig. S7A). The nucleoplasmic and cytoplasmic signals were consistent with the localization of AAR2 and HYL1 in both subcellular compartments (*SI Appendix* Fig. S6;(43)). However, the D-body BiFC signals were surprising as AAR2-YFP did not from nuclear foci in the *AAR2::AAR2-YFP* transgenic line. Over expression with the 35S promoter probably contributed to the signals, but the absence of D-body BiFC signals between AAR2 and DCL1 or AAR2 and SE suggested that interactions between AAR2 and HYL1 occurs in D-bodies, perhaps transiently. This is consistent with the finding that the *aar2-1* mutation affects the formation of HYL1-labeled D-bodies. In a complementary experiment, crude extracts from *AAR2::AAR2-YFP* transgenic plants were subjected to size exclusion chromatography. AAR2-YFP was found to partially co-fractionate with DCL1, HYL1, and SE in high molecular weight fractions (*SI Appendix* Fig. S7B). Together, the results are consistent with the association of AAR2 with the microprocessor in the nucleoplasm. The reduced D-body localization of HYL1 and the positive BiFC signals between AAR2 and HYL1 in D-bodies are consistent with the role of AAR2 in enabling the D-body localization of HYL1. However, the interactions between AAR2 and the microprocessor components were likely weak or transient, as we were unable to detect the interactions by co-immunoprecipitation (*SI Appendix* Fig. S8).

**Fig. 2.**
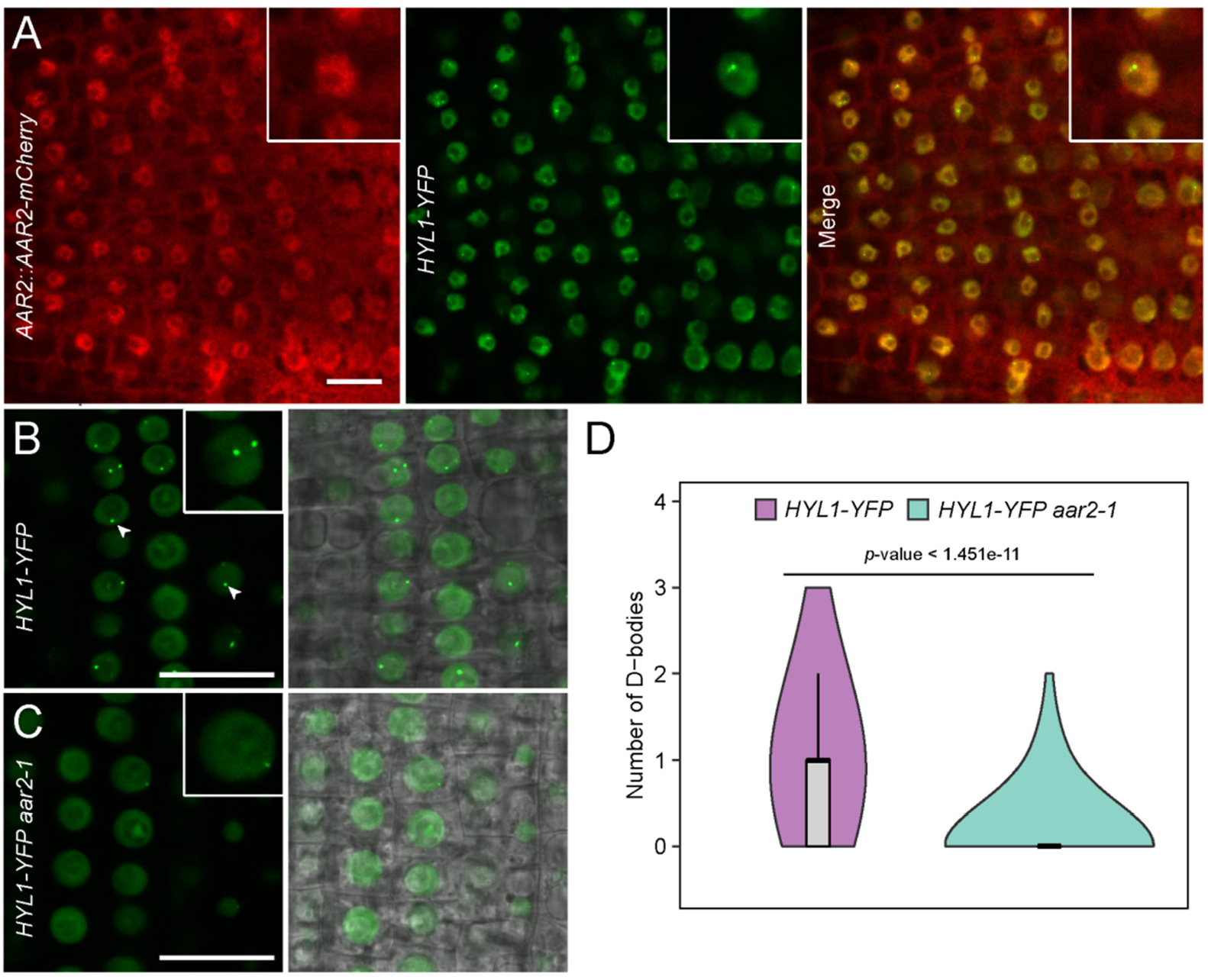
AAR2 associates with microprocessor components and is required for the localization of HYL1 in D-bodies. (*A*) Confocal images of root tip cells expressing both HYL1-YFP and AAR2-mCherry. Enlarged nuclei are shown in the insets. (*B* and *C*) Representative images of HYL1-YFP in WT (*B*) and *aar2-1* (*C*). Arrowheads indicate D-bodies. Scale bar in (*A*-*C*), 20 μm. (*D*) Violin plots showing the number of HYL1-YFP-labeled D-body per cell in WT and *aar2-1*. Quantification was performed by counting more than 400 cells from 15 roots for each genotype. *p*-value was calculated by the Wilcoxon test.

### AAR2 promotes pri-miRNA accumulation without affecting MIR promoter activities

To further investigate how *AAR2* promotes miRNA biogenesis, we examined the levels of pri-miRNAs using qRT-PCR and found that levels of all 16 examined pri-miRNAs except miR167b showed a significant reduction in both *aar2* mutants (Fig. 3A). This indicates that *AAR2* is required for pri-miRNA accumulation. As steady-state pri-miRNA levels are a result of *MIR* transcription, pri-miRNA processing, and pri-miRNA degradation, we first examined whether *MIR* promoter activities were affected in *aar2-1. pMIR167a::GUS*, a GUS reporter for *MIR167a* promoter activity (44), was crossed into *aar2-1*. GUS histochemical staining showed no obvious difference in terms of GUS activity between *aar2-1* and wild type (Fig. 3B). Moreover, qRT-PCR analysis showed that pri-miR167a was significantly reduced in *aar2-1* while GUS transcript levels were comparable between *aar2-1* and wild type (Fig. 3C). We further examined four other *MIR* promoter activity reporters, *MIR165a::GFPer, MIR165b::GFPer, MIR166a::GFPer*, and *MIR166b::GFPer* (45), and found no detectable difference between *aar2-1* and WT in terms of GFP fluorescence intensities or GFP transcript levels, while levels of endogenous pri-miR165A, pri-miR165B, pri-miR166A and pri-miR166B were significantly reduced in *aar2* as compared to wild type (Fig. 3 D-E). Taken together, these results suggest that *AAR2* is required for pri-miRNA accumulation without affecting the transcription of the corresponding *MIR* genes.

**Fig. 3.**
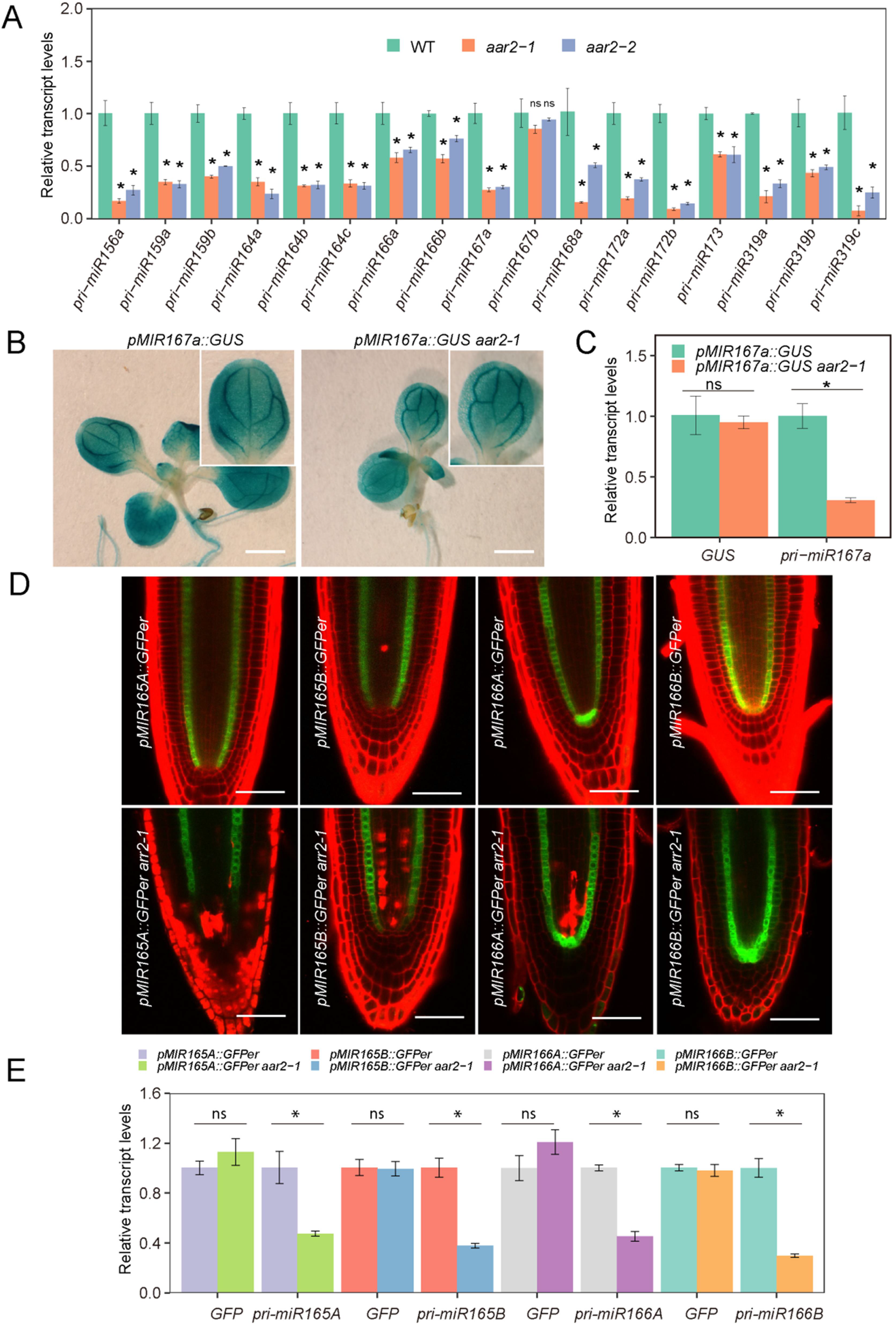
*AAR2* promotes pri-miRNA accumulation without affecting *MIR* promoter activities. (*A*) qRT-PCR analysis of pri-miRNAs in WT, *aar2-1* and *aar2-2. UBQ5* was used as an internal control. (*B*) Representative images of histochemical staining of GUS in *pMIR167a::GUS* and *pMIR167a::GUS aar2-1* transgenic plants. Fifteen plants for each genotype were analyzed. Scale bar, 1 mm. (*C*) Levels of the GUS transcript and pri-miR167a in *pMIR167a::GUS* and *pMIR167a:GUS aar2-1* plants as determined by qRT-PCR. (*D*) Representative images of roots of *MIR165A::GFPer, MIR165B::GFPer, MIR166A::GFPer*, and *MIR166B::GFPer* in WT (top panel) and *aar2-1* (bottom panel). 15 roots for each genotype were analyzed. (*E*) Levels of the GFP transcript and the corresponding endogenous pri-miRNAs in the aforementioned promoter reporter lines as determined by qRT-PCR. Error bars in (*A, C* and *E*) represent SD calculated from three independent replicates. Asterisks indicate significant difference (Student’s t-test, *P < 0.05). ns, no significant difference.

### Mutations in AAR2 lead to pri-miRNA degradation in a HYL1-dependent manner

The association of AAR2 with the microprocessor and its requirement for the formation of HYL1-labeled D-bodies prompted us to test the genetic interactions between *aar2-1* and mutations in *HYL1* and *SE*. The *se-1 aar2-1* double mutant showed more severe developmental phenotypes than either the *aar2-1* or *se-1* single mutant, while the *hyl1-2 aar2-1* double mutant largely resembled *hyl1-2* in plant morphology (Fig. 4A). Consistent with these observations, miRNA levels in the *se-1 aar2-1* double mutant were further decreased in comparison with those in either the *aar2-1* or *se-1* single mutant while miRNA levels in the *hyl1-2 aar2-1* double mutant were comparable to those in *hyl1-2* (Fig. 4B), suggesting that *AAR2* and *SE* contribute additively to miRNA biogenesis while *AAR2* contributes to miRNA biogenesis through the *HYL1*.

**Fig. 4.**
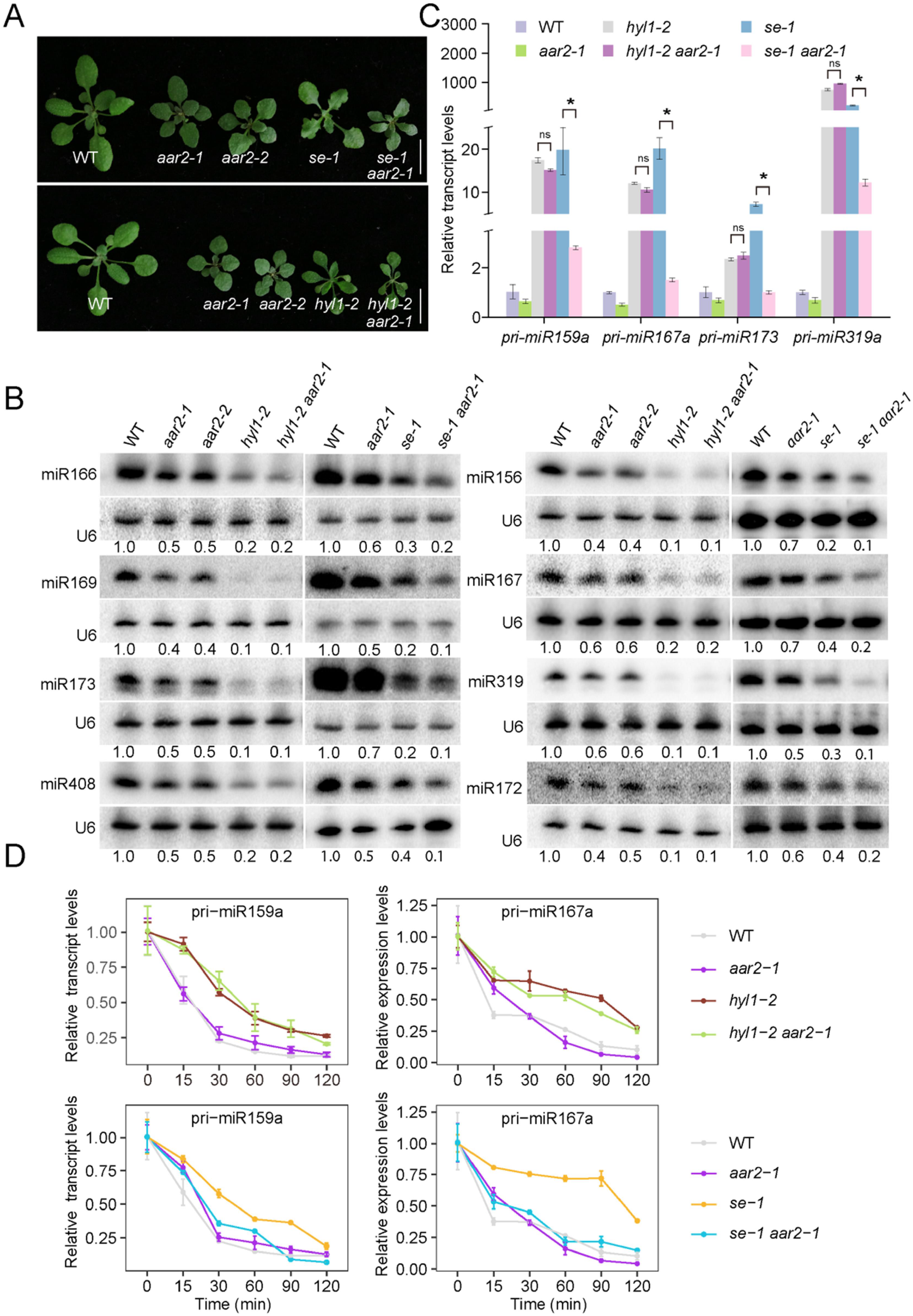
AAR2 affects miRNA biogenesis in a HYL1-, but not SE-dependent manner. (*A*) Phenotypes of 3-week-old plants of the indicated genotypes. Scale bar, 10 mm. (*B*) RNA gel blot analysis of miRNA accumulation in *se-1 aar2-1* and *hyl1-2 aar2-1* double mutants as compared to the corresponding single mutants. U6 was used as the loading control. The numbers below the blots represent relative miRNA levels. (*C*) qRT-PCR analysis of pri-miRNAs in *hyl1-2 aar2-1* and *se-1 aar2-1* as compared to the corresponding single mutants. *UBQ5* was used as an internal control. Error bars represent SD calculated from three independent replicates. Asterisks indicate significant difference (Student’s t-test, *P < 0.05). (*D*) Analysis of pri-miRNA decay upon treatment of WT, *aar2-1, hyl1-2, se-1, hyl1-2 aar2-1* and *se-1 aar2-1* seedlings with the transcription inhibitor cordycepin. *UBQ5* was used as an internal control. Error bars indicate SD from three technical replicates.

The *aar2-1* mutation caused a reduction in the levels of pri-miRNAs without affecting *MIR* gene transcription. We investigated whether this reduction of pri-miRNA accumulation still occurred in the *hyl1* or *se* background, in which pri-miRNAs accumulate to higher levels due to inefficient processing (46). Levels of examined pri-miRNAs were reduced in the *se-1 aar2-1* double mutant relative to *se-1*, but were similar in the *hyl1-2 aar2-1* double mutant as compared to *hyl1-2* (Fig. 4C), which indicates that the effects of *aar2-1* on pri-miRNA accumulation were independent of *SE* but dependent on *HYL1*.

To determine how *aar2-1* results in lower pri-miRNA levels, we observed the decay of pri-miRNAs in seedlings treated with the transcription inhibitor cordycepin. When transcription is inhibited, the levels of pri-miRNAs are expected to decrease due to DCL1 processing as well as RNA degradation. Consistent with findings from a previous study (35), in both *se-1* and *hyl1-2*, pri-miRNA levels decreased more slowly than in wild type, probably due to the lack of pri-miRNA processing (Fig. 4D). The half-lives of examined pri-miRNAs were similar between *aar2-1* and wild type (Fig. 4D). However, the half-lives of pri-miRNAs were shorter in the *se-1 aar2-1* double mutant as compared to *se-1* (Fig. 4D), suggesting that the *aar2-1* mutation led to the degradation of pri-miRNAs. Intriguingly, the half-lives of pri-miRNAs were similar in the *hyl1-2 aar2-1* double mutant and the *hyl1-2* single mutant, suggesting that the accelerated pri-miRNA degradation caused by *aar2-1* is dependent on HYL1.

### AAR2 is involved in HYL1 degradation

Previous studies showed that HYL1 is an unstable protein with relatively short half-lives particularly under darkness (31, 47). That *AAR2* affects miRNA biogenesis through *HYL1* prompted us to explore the possibility that *AAR2* plays a role in the regulation of HYL1 protein stability. 10-day-old seedlings were transferred to darkness for 2 days and HYL1 protein levels were determined in wild type and *aar2* mutants. Under dark treatment, HYL1 levels decreased in wild type, which is consistent with previous findings (31). HYL1 protein levels were higher in *aar2-1* and *aar2-2* relative to wild type after dark treatment (Fig. 5A). Treatment of seedlings with carbobenzoxy-Leu-Leu-leucinal (MG132) while they were in darkness led to higher levels of HYL1 both in wild type and *aar2* (Fig. 5B). These results indicate that the degradation of HYL1 is impaired in *aar2*. To test whether AAR2 affects HYL1 degradation under the light condition, we performed a protein decay assay by treating long-day grown, 10-day-old seedlings with cycloheximide (CHX) during the light period to block de novo protein synthesis and examined HYL1 protein levels at 2 hours after treatment. The levels of HYL1 showed a 50% decrease after 2 hours of CHX treatment in wild type but the decay rate was much slower in *aar2* mutants (Fig. 5C). Moreover, MG132 treatments notably suppressed the degradation of HYL1 in both wild type and *aar2* (Fig. 5D). Therefore, AAR2 is required for the degradation of HYL1.

**Fig. 5.**
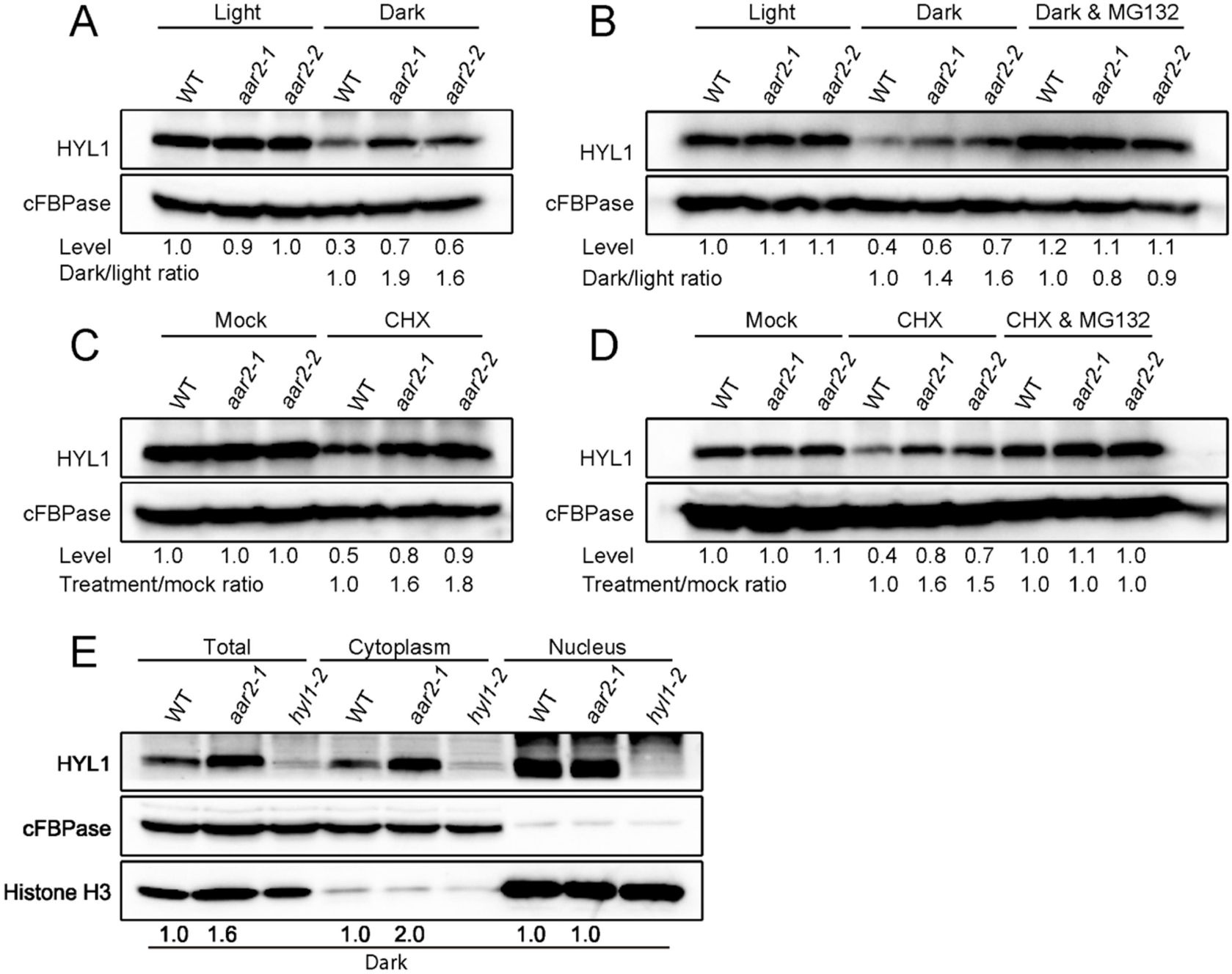
AAR2 affects HYL1 degradation. (*A-B*) Protein gel blot analysis of HYL1 in WT, *aar2-1* and *aar2-2* seedlings kept in the light or transferred to the dark and grown for 2 days (*A*) or incubated with MG132 during the dark period (*B*). HYL1 levels were normalized against cFBPase and expressed in values relative to WT in the light. The dark/light ratio represents the ratio of HYL1 levels in the dark and in the light and is relative to WT. (*C-D*) Protein gel blot analysis of HYL1 in long-day grown WT, *aar2-1* and *aar2-2* seedlings treated with cycloheximide (CHX) (C) or CHX together with MG132 (D) for 2 h. HYL1 levels were normalized against cFBPase and expressed in values relative to the WT mock-treated samples. The treatment/mock ratio represents the ratio of HYL1 levels in mock-treated samples and in samples treated with CHX or CHX+MG132 and is relative to WT. (*E*) HYL1 levels in total, cytoplasmic and nuclear fractions under dark treatment as described in (*A*). cFBPase and histone H3 were used as cytoplasmic and nuclear markers, respectively. HYL1 levels were normalized against cFBPase or histone H3 and expressed in values relative to WT.

As HYL1 is degraded in the cytoplasm under darkness (31), we performed nuclear-cytoplasmic fractionation to examine the distribution of HYL1 in the dark. Results showed that HYL1 levels were increased in *aar2-1* specifically in the cytoplasm as compared to wild type (Fig. 5E). Moreover, there was no obvious difference in HYL1 levels between *aar2-1* and wild type in either total, cytoplasmic or nuclear fractions under the light conditions (*SI Appendix* Fig. S9A), suggesting that *aar2-1* does not affect HYL1’s nuclear-cytoplasmic partitioning. Similarly, RNA gel blot analysis showed that *aar2-1* did not alter the nucleocytoplasmic distribution of miRNAs (*SI Appendix* Fig. S9B). Taken together, these results support that AAR2 is involved in HYL1 degradation in the cytoplasm but does not affect miRNA or HYL1’s nuclear-cytoplasmic distribution.

### AAR2 is required for HYL1 dephosphorylation

Phosphorylated HYL1 is inactive in miRNA biogenesis but forms a reserve pool of HYL1 that is resistant to degradation (32). As HYL1 is more resistant to degradation in *aar2* mutants, we thought that the phosphorylation status of HYL1 might be affected in the mutants. CPL1 is a phosphatase known to dephosphorylate HYL1 (28). We introduced the *cpl1-7* mutation into *aar2-1* and found that the *aar2-1 cpl1-7* double mutant showed more severe developmental phenotypes than either *aar2-1* or *cpl1-7* (*SI Appendix* Fig. S10A). The phosphorylation status of HYL1 was analyzed using phos-tag gels and independent experiments showed that the hypo-phosphorylated form of HYL1 was reduced in *aar2-1* as compared to wild type (Fig. 6A and *SI Appendix* Fig. S10B). We examined pri-miRNA accumulation in the *aar2-1 cpl1-7* double mutant using qRT-PCR. Results showed that levels of all examined pri-miRNAs were decreased in both *aar2-1* and *cpl1-7*, while a further decrease was observed for some pri-miRNAs in the *aar2-1 cpl1-7* double mutant as compared to either *aar2-1* or *cpl-17* (*SI Appendix* Fig. S10C). Moreover, the levels of mature miRNAs were reduced in the *aar2-1 cpl1-7* double mutant as compared to either single mutant (*SI Appendix* Fig. S10D).

**Fig. 6.**
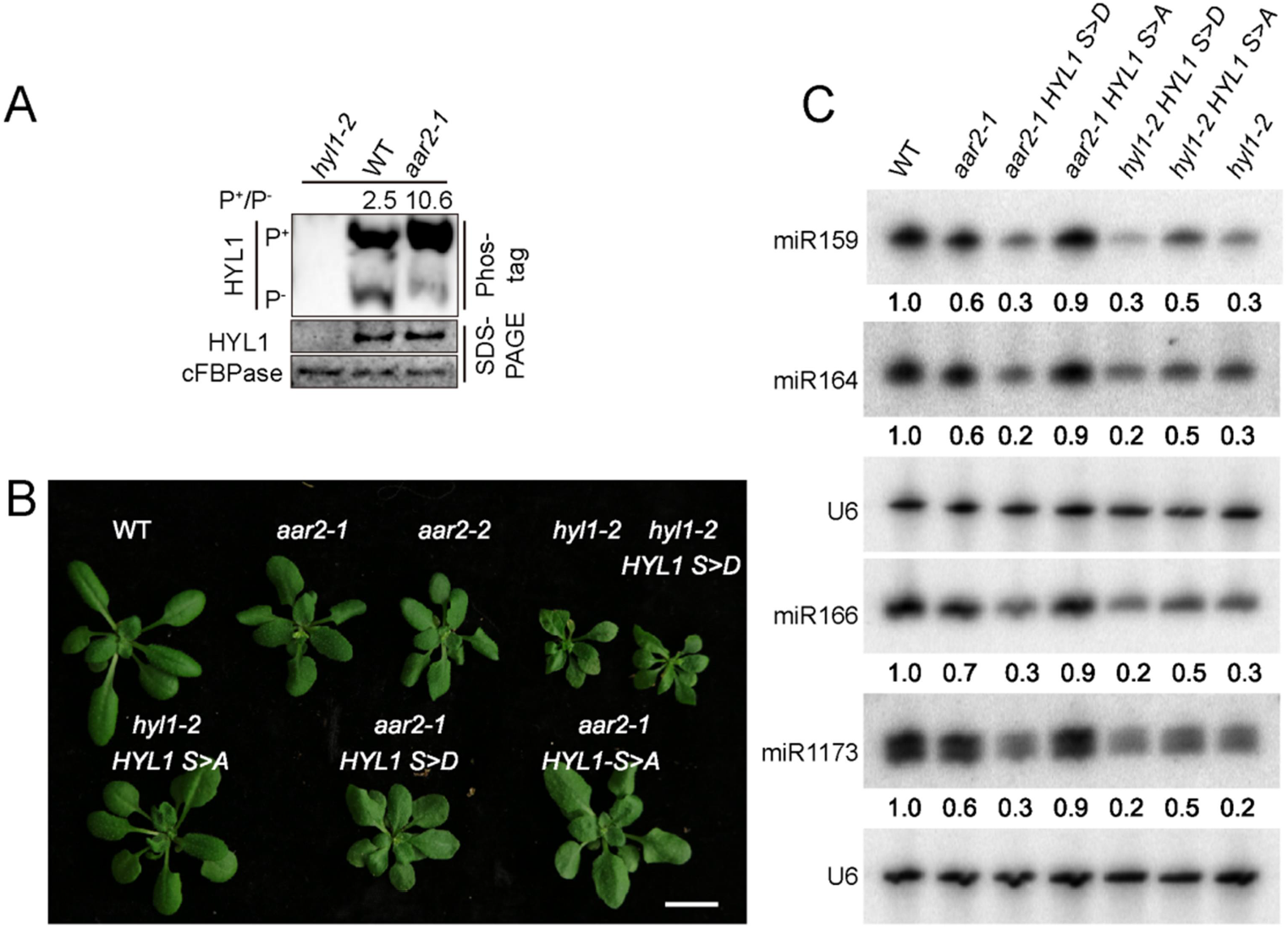
AAR2 affects HYL1 phosphorylation status. (*A*) Phosphoprotein mobility shift gel detecting phosphorylated (P^+^) and non-phosphorylated (P^-^) HYL1 in the indicated genotypes. SDS-PAGE gels in the middle and bottom panels show the levels of HYL1 and cFBPase (used as a loading control), respectively. (*B*) 3-week-old plants of the indicated genotypes. Scale bar, 10 mm. (*C*) RNA gel blot analysis of miRNA accumulation in the indicated genotypes. U6 was used as the loading control. The numbers represent relative abundances.

We next tested whether *AAR2* affects miRNA biogenesis through HYL1 dephosphorylation. Expression of HYL1 S>A, a non-phosphorylatable HYL1 with all seven predicted serine phosphorylation sites mutated to alanine (28), partially rescued the developmental defects of *aar2*, while *aar2-1* plants expressing HYL1 S>D, a phosphomimetic HYL1 with all seven predicted serine phosphorylation sites mutated to aspartic acid (28), showed more severe developmental defects (Fig. 6B). We examined miRNA accumulation in HYL1 S>D *aar2-1* and HYL1 S>A *aar2-1* using RNA gel blotting. As expected, the accumulation of all examined miRNAs was partially rescued in *hyl1-2* by HYL1 S>A while HYL1 S>D was unable to rescue the miRNA biogenesis defects in *hyl1-2* (Fig. 6C and *SI Appendix* Fig. S10E). Accordingly, pri-miRNA levels were unchanged in *hyl1-2 HYL1 S>D* and reduced in *hyl1-2 HYL1 S>A* relative to *hyl1-2* (*SI Appendix* Fig. S10F). The accumulation of some of the examined miRNAs was also partially rescued in *aar2-1* by HYL1 S>A while HYL1 S>D led to a further reduction in the levels of all examined miRNAs in *aar2-1* (Fig. 6C and *SI Appendix* Fig. S10E). These results indicate that AAR2 promotes miRNA accumulation through regulating the HYL1 phosphorylation status. The accumulation of phosphorylated HYL1 in *aar2* mutants (Fig. 6A and *SI Appendix* Fig. S10B) could also explain the increased stability of this protein in the mutants.

### AAR2 is required for efficient messenger RNA splicing

Homologs of AAR2 in yeast and mammals are involved in the assembly of U5 snRNP, an essential complex in pre-mRNA splicing (38, 48). In *Arabidopsis*, a role of AAR2 in splicing has never been characterized. This prompted us to examine whether *aar2* mutations show splicing defects. We performed RNA-seq in WT, *aar2-1* and *aar2-2* with three biological replicates. In total, we identified 1162 up-regulated genes (hyper DEGs) and 1078 down-regulated genes (hypo DEGs) in *aar2-1*, and 647 hyper DEGs and 924 hypo DEGs in *aar2-2*. 459 up-regulated genes and 604 down-regulated genes were found in both *aar2-1* and *aar2-2* (*SI Appendix* Fig. S11A and Dataset S2). Gene Ontology (GO) enrichment analysis showed that both common hyper-DEGs and hypo-DEGs were enriched in genes involved in responses to stimuli (*SI Appendix* Fig. S11B).

Analyses of the RNA-seq data revealed that both *aar2-1* and *aar2-2* show global splicing defects with 512 and 1159 intron retention events in *aar2-1* and *aar2-2*, respectively. Two examples (AT4G35760 and AT3G56590) are presented (Fig. 7A and B and Dataset S3). *HYL1* transcript level or intron splicing was not affected in *aar2-1* or *aar2-2*, which further confirmed that the role of AAR2 in miRNA biogenesis is not through modulating *HYL1* expression (*SI Appendix* Fig. S12 A-B). We further compared the retained introns between *aar2* and *mac3a mac3b* or *prl1 prl2*, which are mutations in the MAC complex required for both miRNA biogenesis and pre-mRNA splicing (49). No overlap was found between retained introns in *aar2* and either *mac3a mac3b* or *prl1 prl2* (Fig. 7C), suggesting that AAR2 and MAC act on different sets of introns. Next, we examined whether the genes with retained introns in *aar2* had any common features. When compared to all genes, the genes with intron retention defects in *aar2* tended to have more introns (Fig. 7D).

**Fig. 7.**
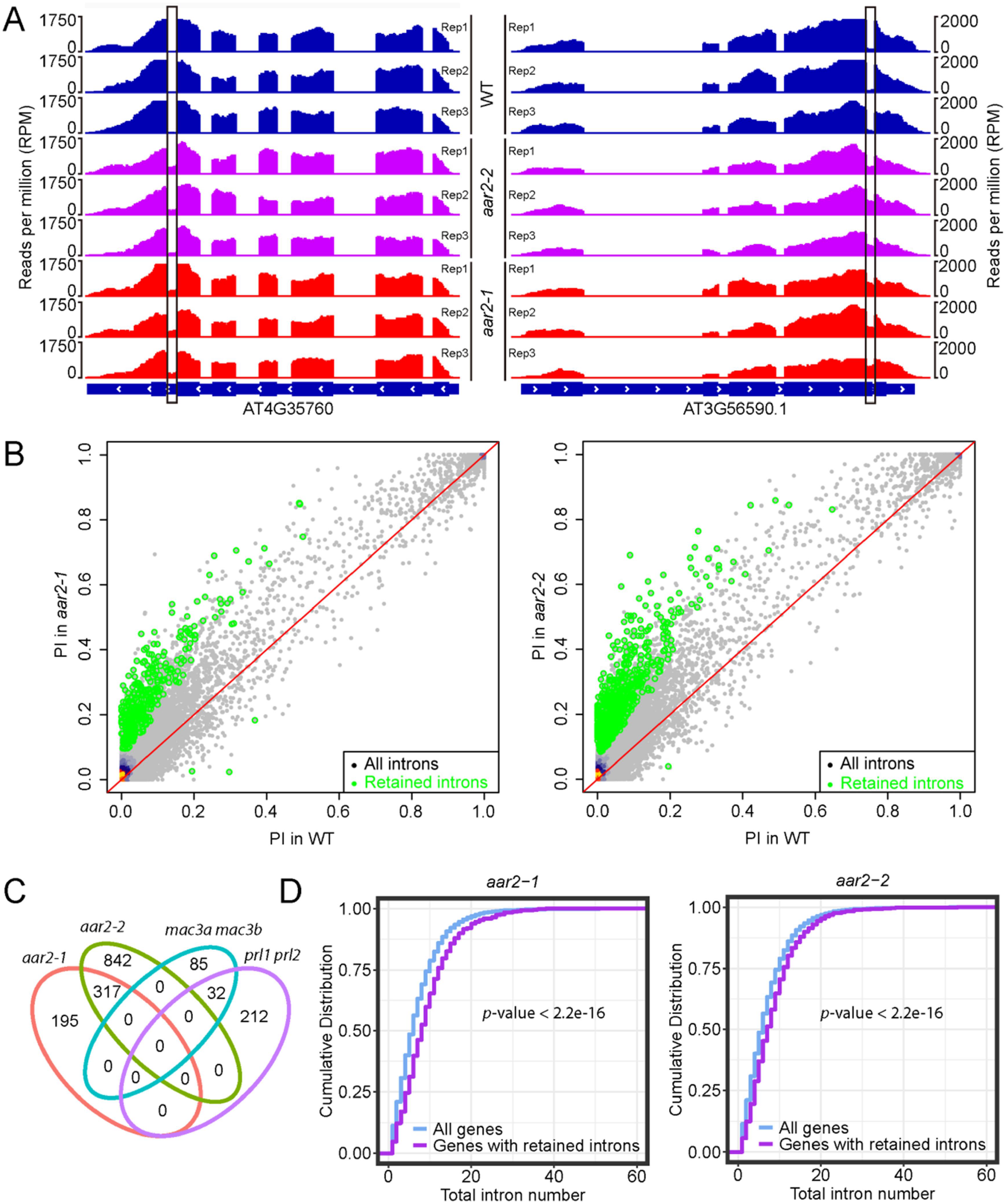
AAR2 is required for efficient pre-mRNA splicing. (*A*) Examples of two genes with intron retention defects in the *aar2-1* and *aar2-2* mutants. RNA-seq reads are shown against the gene models below. In the gene models, exons and introns are marked with rectangles and lines, respectively. The retained introns in the *aar2-1* and *aar2-2* mutants are marked with black rectangles. (*B*) Scatter plots showing the percentage of retained introns (PI) in *aar2-1* and *aar2-2* mutants. The green dots represent introns with statistically significant retention defects in the mutants. (*C*) Venn diagrams showing the number of retained introns in *aar2-1, aar2-2, mac3a mc3b* and *prl1 prl2* mutants, and the overlap of the introns retained among these mutants. (*D*) Cumulative density plots of intron number in all genes and genes with retained introns in the *aar2-1* and *aar2-2* mutants. *p*-values were calculated by the Wilcoxon test.

## Discussion

AAR2 is a conserved protein in eukaryotes. Its homologs in yeast and humans globally impact pre-mRNA splicing through a role in U5 snRNP assembly, but AAR2’s function in plants has never been characterized. In this study, we show that *Arabidopsis* AAR2 indeed plays a global role in splicing, with hundreds to thousands of introns being retained in *aar2* mutants. We show that Arabidopsis AAR2 is required for miRNA accumulation in addition to its conserved function in splicing. In fact, a global reduction in the levels of 24-nt small RNAs was also found in the *aar2-1* mutant, suggesting that AAR2 also promotes siRNA accumulation. Several other splicing factors were also previously shown to impact both miRNA and siRNA accumulation (20, 44, 50), which indicates that mutations in these genes may affect the splicing of a gene(s) involved in siRNA biogenesis and indirectly affect siRNA accumulation. However, it is also possible that these genes are directly involved in Pol IV/RDR2-dependent siRNA biogenesis. We demonstrate that AAR2 promotes miRNA biogenesis through regulating the phosphorylation status of the microprocessor component HYL1. As miRNA biogenesis requires double-stranded RNA binding proteins, such as HYL1, TRBP, and DGCR8, in both plants and animals, it is possible that animal AAR2 exerts a similar effect on miRNA biogenesis.

HYL1 is a short-lived protein and is degraded by the cytoplasmic protease HYL1-CLEAVAGE SUBTILASE 1 (HCS1) in a light-dependent manner (51). Under light conditions, cytoplasmic CONSTITUTIVE PHOTOMORPHOGENIC 1 (COP1) suppresses the activity of HCS1 to prevent HYL1 from degradation, while under dark conditions, the relocation of COP1 to the nucleus releases its repression on HCS1, leading to HYL1 degradation (31, 47, 51, 52). We found that the rate of HYL1 degradation in the dark is slower in *aar2* than WT. Furthermore, HYL1 accumulated significantly in the cytoplasmic fraction in *aar2* under dark treatment, which is consistent with the current model that HYL1 is degraded in the cytoplasm and suggests that AAR2 is required for HYL1 degradation in the cytoplasm.

The degradation of HYL1 is impacted by its phosphorylation status. Hyper-phosphorylated HYL1 tends to be sequestered in the nucleus and protected from degradation in the cytoplasm in the dark (32). The reduced rate of HYL1 degradation in *aar2* mutants in the dark prompted us to examine the phosphorylation status of HYL1 in *aar2* mutants. Our results showed that the ratio of non-phosphorylated to phosphorylated HYL1 is significantly lower in *aar2* even in seedlings grown under long-day conditions. The non-phosphorylated form of HYL1 is the active form in miRNA biogenesis and tends to form D-bodies in the nucleus (28). Thus, the reduction in D-body number and in miRNA accumulation in *aar2* mutants could be explained by defects in HYL1 dephosphorylation. Consistently, expression of non-phosphorylatable HYL1 partially rescues the miRNA biogenesis defects of *aar2-1*. Taken together, the findings from this study suggest that AAR2 promotes miRNA accumulation through HYL1 dephosphorylation, stability, and localization to D-bodies (*SI Appendix* Fig. S13).

This study also revealed a previously unknown role of HYL1 in the degradation of pri-miRNAs. In mutants in core microprocessor components, such as *dcl1, hyl1* and *se*, pri-miRNAs accumulate to higher levels as compared to wild type as a consequence of compromised processing. In *aar2* mutants, despite a decrease in the active form of HYL1 and a presumed decrease in pri-miRNA processing, pri-miRNA levels are lower than in wild type. Moreover, the promoter activities of five examined *MIR* genes were unaffected in *aar2-1* while the corresponding pri-miRNAs were decreased in abundance, which indicates that *aar2-1* led to reduced pri-miRNA stability. After *MIR* gene transcription, both processing by the microprocessor and degradation are expected to contribute to the levels of pri-miRNAs in vivo. In order to examine the effects of *aar2-1* on pri-miRNA degradation, we followed the decrease in pri-miRNA levels in the *se-1* background upon inhibition of transcription to exclude the effects of pri-miRNA processing. Indeed, the *aar2-1* mutation led to faster pri-miRNA degradation in the *se-1* background. It should be noted that although pri-miRNA levels were reduced in *aar2*, pri-miRNAs decreased similarly in *aar2* and wild type in the time course of cordycepin treatment (Fig 4D). One possible explanation is that the processing efficiency of pri-miRNAs is lower in *aar2* as compared with WT due to the reduced levels of non-phosphorylated HYL1, and the reduced processing would increase the half-life of pri-miRNAs in the assay. Most intriguingly, pri-miRNA levels were similar in the *hyl1-2 aar2-1* double mutant and the *hyl1-2* single mutant (Fig 4C), and consistently, the decay of pri-miRNAs was similar in *hyl1-2 aar2-1* and *hyl1-2* in the time course of cordycepin treatment (Fig 4D). We conclude that AAR2 protects pri-miRNAs from HYL1-dependent degradation. This finding, together with those from another study (35), reveal that HYL1 and SE not only promote pri-miRNA processing but also help degrade pri-miRNAs. Perhaps they exert these seemingly opposite functions through distinct interaction partners or at different subcellular locations. Intriguingly, a previous study showed that HYL1 protects pri-miRNAs from degradation by the nuclear exosome (53). Therefore, HYL1 may play opposite roles in pri-miRNA stability under different conditions or in different forms. As *aar2* mutants accumulate phosphorylated HYL1, we hypothesized that this isoform, so far considered inactive, may have a specific function in promoting pri-miRNA degradation. However, expression of HYL1 S>D in *hyl1-2* did not lead to a reduction in pri-miRNA levels (*SI Appendix* Fig. S10F).

Pre-mRNA splicing and miRNA biogenesis can be two independent processes (54). However, several proteins that have been shown to be involved in pre-mRNA splicing also facilitate miRNA biogenesis (18-20, 49). Yeast and mammalian AAR2 is a conserved U5 snRNP assembly factor, serving as a component of the cytoplasmic U5 snRNP precursor. Upon importing into the nucleus, AAR2 is phosphorylated and replaced by Brr2 to promote the maturation of U5 snRNP (38, 55). We showed that Arabidopsis AAR2 is localized in both the cytoplasm and the nucleus, and is required for the splicing of hundreds to thousands of introns. Thus, it is possible that Arabidopsis AAR2 functions in U5 snRNP assembly. In Arabidopsis, only a subset of pri-miRNAs harbors introns (56, 57), however, miRNA accumulation is non-discriminately affected in *aar2* mutants regardless of whether the pri-miRNAs contain introns or not, which indicates that the miRNA biogenesis function of AAR2 is independent of its function in splicing.

## Methods

### Plant materials and growth conditions

All *Arabidopsis thaliana* strains used in this study are in the Columbia-0 accession. The *SUC2::atasi-SUL* transgenic line used in the mutagenesis screen is a kind gift from Dr. Detlef Weigel (37). *aar2-1* is a new allele isolated from the *SUC2::atasi-SUL* mutagenesis screening. The T-DNA insertion lines *aar2-2* (SALK SALK_117746) and *hyl1-2* (SALK_064863) were obtained from ABRC. *se-1* and *cpl1-7* were previously described (28, 58). 35S::HYL1 S>D (7 serine codons (S7, S8, S42, S60, S85, S89, S159) mutated to aspartate) and 35S::HYL1 S>A 7 serine codons (S7, S8, S42, S60, S85, S89, S159) mutated to alanine) transgenic lines were previously described (32). To generate *HYL1-YFP aar2-1, pMIR167a::GUS aar2-1, MIR165A::GFPer aar2-1, MIR165B::GFPer aar2-1, MIR166A::GFPer aar2-1* and *MIR166A::GFPer aar2-1, aar2-1* was crossed with *HYL1-YFP* (59), *pMIR167a::GUS* (44), *MIR165A::GFPer, MIR165B::GFPer, MIR166A::GFPer* and *MIR166A::GFPer* (45), respectively. All primers used for genotyping are listed in Dataset S4.

Seeds were sown on ½ Murashige and Skoog Basal medium containing 1% sucrose and plants were grown under long-day conditions (16 h light and 8 h dark) at 22°C. The plants were either collected at 14 days for analysis or transferred to soil for phenotype observation. For transient dark treatment, 10-day-old seedlings were grown under long-day conditions and transferred to darkness for 48 hours.

### Mutagenesis screening and AAR2 identification

The *SUC2::atasi-SUL* transgenic lines were described previously (37). A transgenic line with a single-locus T-DNA insertion was used for the ethyl methansulfonate mutagenesis as described (37). A mutant with reduced leaf yellowing was isolated and back-crossed with the parental line *SUC2::atasi-SUL*. In the F2 population, ∼100 plants with the *SUC2::atasi-SUL aar2-1* phenotype were collected and genomic DNA was extracted and subjected to library construction. The library was pair-end sequenced on the Illumina platform Hiseq2000 at 50X coverage. A mutation in AT1G66510 (*AAR2*) was identified and further validated using Derived Cleaved Amplified Polymorphic Sequences (dCAPS) analysis in the F2 population. Primers for the dCAPS analysis are listed in Dataset S4.

### DNA constructs and plant transformation

A 5.2 kb genomic fragment of *AAR2* containing the promoter and coding region without the stop codon was PCR-amplified and cloned into pENTR/D-TOPO (Invitrogen) and then introduced into pGWB40 (60) to generate *AAR2::AAR2-YFP*. For the *AAR2::AAR2-mCherry* construct, the mCherry coding sequence and the *AAR2* genomic fragment were PCR-amplified and recombined into the pCambia1300 vector (abcam, ab275754). For *35S::AAR2-nYFP, AAR2* cDNA was PCR-amplified and inserted downstream of the 35S promoter and fused with nYFP in pXY103 (61). The above constructs were introduced into *aar2-1* by the *Agrobacterium tumefaciens*-mediated floral dipping method (62). The primers used are listed in Dataset S4.

### Small RNA gel blot analysis

Total RNAs were extracted from 14-day-old seedlings with TRI reagent (Molecular Research Center). Small RNA gel blot analysis was performed as described previously (63). 5-10 μg total RNA was separated in a 15% Urea-PAGE gel and transferred onto a Hybond NX membrane. The membrane was probed by 5’-end ^32^P-labled anti-sense DNA oligonucleotides to detect miRNAs or ta-siRNAs. The relative levels of miRNAs or ta-si-RNAs were calculated by normalizing against the internal control U6 or tRNA.

### Quantitative RT-PCR

Total RNAs were treated with DNase I (Roche) followed by cDNA synthesis using RevertAid reverse transcriptase (Thermo Fisher Scientific) with oligo(dT) primers. Quantitative RT-PCR was performed using iQ SYBRGreen Supermix (BioRad) on the BioRad CFX96 system. Primers used are listed in Dataset S4.

### RNA stability measurements

RNA stability measurements were carried out as described (64). Briefly, 12-day-old seedlings were transferred from ½ MS medium agar plates to 12-well plates with 2 ml ½ MS liquid medium and incubated overnight to equilibrate the seedlings. Cordycepin was added to a final concentration of 0.6 mM and seedlings were collected at various time points (0 min, 15 min, 30 min, 60 min, 90 min, 120 min) for RNA extraction. Quantitative RT-PCR was performed to determine the levels of pri-miRNAs.

### Cycloheximide (CHX) and MG132 treatments

CHX treatment of *Arabidopsis* seedlings were performed as previously described (31). 10-day-old seedlings were treated with 0.5 mM CHX with or without 50 μM MG132 for 2 h. Samples were collected and subjected to protein gel blot analysis to determine HYL1 protein levels.

### Microscopy and imaging

For Gus staining, whole seedlings from WT and *aar2-1* with a homozygous *pMIR167a::GUS* transgene were incubated with the GUS staining solution (65) at 37°C in the dark. Tissues were cleared with 70% ethanol before being observed under a stereomicroscope (Leica).

For propidium iodide staining, seedlings were incubated with 10 μM propidium iodide for 10 min and then washed with distilled water. After staining, the roots were excised, mounted in water and observed under a confocal microscope.

### Gel filtration assay

0.5 g 14-day-old *AAR2::AAR2-YFP* seedlings was ground into fine powder. 1.5ml 1XPBS supplemented with 1x protease inhibitor cocktail (Roche) was added into the powder and the suspension was incubated at 4°C for 30 min. The suspension was centrifuged at 4°C for 20 min at 13200g and the supernatant was transferred into a new tube. Upon a second centrifugation, the supernatant was collected and filtered through a 0.45 μm filter. Then, 500 μl samples were loaded onto a Superdex 200 Increase 10/300 GL column. 22 fractions were collected and used for protein detection by protein gel blot analysis.

### RNA and small RNA sequencing

For small RNA library construction, 50 μg total RNAs were resolved in a 15% Urea-PAGE gel and 15-40 nt small RNAs were excised from the gel and used as the input for library construction. The libraries from various genotypes and/or replicates were pooled and sequenced on an Illumina HiSeq 2500 platform. Small RNA-seq data were analyzed with the pRNASeqTools pipeline (https://github.com/grubbybio/pRNASeqTools). The raw reads were first trimmed to remove the adapter sequence (AGATCGGAAGAGC) using Cutadapt3.0 (66). The trimmed reads were mapped onto the *Arabidopsis thaliana* genome Araport11 using Shortstack (67) with parameters ‘-bowtie_m 1000 -ranmax 50 -mmap u -mismatches 0’.

Normalization was performed by calculating the RPMR (reads per million of 45S rRNA reads) value. Differentially accumulated small RNAs were analyzed using the R package DESeq2 (68).

For RNA-seq library construction, poly(A) RNAs were isolated from total RNAs using NEBNext Poly(A) mRNA Magnetic Isolation Module. RNA-seq libraries were prepared using purified poly(A) RNA with NEBNext® Ultra™ II RNA Library Prep Kit for Illumina and sequenced on an Illumina HiSeq 2500 platform. Data were analyzed using the pRNASeqTools pipeline (https://github.com/grubbybio/RNASeqTools). Briefly, reads were mapped to the *Arabidopsis* genome Araport11 using STAT v2.6(69) with default settings. Mapped reads were counted using featureCounts v1.64(70) and differentially expressed genes were analyzed using the R package DESeq2. The analysis of intron retention in *aar2* and WT was performed using SQUID (https://github.com/sfli001/SQUID).The levels of retained introns were calculated as PI_Junction: (intron-exon junction reads/[intron-exon junction reads + exon-exon reads]).

### Protein gel blot analysis

To determine protein levels, total proteins were extracted from 14-day-old seedlings, resolved in SDS-PAGE gels and then transferred to nitrocellulose membranes. The membranes were blocked with 1x PBS supplemented with 5% (w/v) non-fat milk and 0.05% Tween 20 and then incubated with primary antibodies (Anti-HYL1, 1:2000, Agrisera, AS06136; Anti-cFBPase, 1:5000, Agrisera, AS04043; Anti-Histone H3, 1:3000, abcam, ab1791; anti-SE, 1:2000, Agrisera, AS09532A; anti-GFP, 1:1000, Sigma-Aldrich, 11814460001; Homemade DCL1, 1:2000,) at 4°C. After three washes with 1x PBS supplemented with 0.05% Tween 20, the membranes were incubated with secondary antibodies and then detected with ECL reagent (Amersham, RPN2236). HYL1 phospho-isoforms were detected using Phos-tag SDS-PAGE (FUJIFILM Wako Pure Chemical Corporation, 190-16721) following manufacturer’s instructions. Briefly, total proteins were separated on a Phos-tag SDS-PAGE gel, transferred onto a PVDF membrane using the wet-tank transfer method, and then probed with the HYL1 antibody.

### Nuclear-cytoplasmic fractionation

Nuclear-cytoplasmic fractionation was performed as described (71). 1 g of 14-day-old seedlings was ground into fine powder in liquid nitrogen and the powder was resuspended in 2 ml lysis buffer (20 mM Tris-HCl, pH7.5, 25% glycerol, 250 mM sucrose, 20 mM KCl, 2.5 mM MgCl_2_, 2 mM EDTA, 5 mM DTT and 1x protease inhibitor cocktail). The lysates were filtered through two layers of Miracloth. The flow-through was centrifuged at 1500g at 4°C for 10 min. The supernatant was collected and centrifuged at 10000g at 4°C for 15 min and then the supernatant was kept as the cytoplasmic fraction. The pellet was washed 6 times with nuclear resuspension buffer (20 mM Tris-HCl pH 7.5, 2.5 mM MgCl_2_, 25% glycerol, and 0.2% Triton X-100) and then resuspended with 500 μL NRB2 (20 mM Tris-HCl, pH 7.5, 10 mM MgCl_2_, 250 mM Sucrose, 0.5% Triton X-100, 5 mM β-mercaptoethanol and 1x protease inhibitor cocktail) and carefully overlaid on top of 500 μL NRB3 (20 mM Tris-HCl pH 7.5, 10 mM MgCl_2_, 1.7 M sucrose, 0.5% Triton X-100, 5 mM β-mercaptoethanol and 1x protease inhibitor cocktail), followed by centrifugation at 16000g for 45 min at 4 °C The nuclear and cytoplasmic fractions were subjected for RNA extraction by TRI reagent (Molecular Research Center). For protein gel blot analysis, the nuclear and cytoplasmic fractions were boiled in 1 X SDS sample loading buffer (50 mM Tris-Cl at pH 6.8, 2% SDS, 0.1% bromophenol blue, 10% glycerol, 1% 2-mercaptoethanol) before being resolved in SDS-PAGE gels.

### BiFC analysis

BiFC analysis was carried out as described (72). HYL1 cDNA fused with an N-terminal fragment of YFP and HYL1, DCL1 and SE cDNA fused with a C-terminal fragment of YFP were described (14). The constructs containing the constitutive promoter and gene fusions were transformed into *Agrobacterium* and co-infiltrated into *N. benthamiana* leaves. 48 h after infiltration, YFP signals were detected under confocal microscopy (Leica Sp5).

## Acknowledgements

We would like to thank Dr. Detlef Weigel for sharing the *SUC2::atasi-SUL* transgenic line. This work was funded by NIH GM129373 to X.C.

## Author contributions

L.F., B.G., Z.Y. and X.C. designed research; L.F., B.G., N.P.A., and P.A.M. performed research; L.F., B.G., B.L., C.Y., S.L., and X.C. analyzed data; L.F., B.G., N.F. and X.C. wrote the manuscript.

## Data deposition

Genomics datasets have been deposited in the National Center for Biotechnology Information Gene Expression Omnibus database (accession numbers GSE201940 and GSE201941).

