## Supplemental information for "Arabidopsis AAR2, a conserved splicing factor in eukaryotes, acts in microRNA biogenesis"

### Figure S1

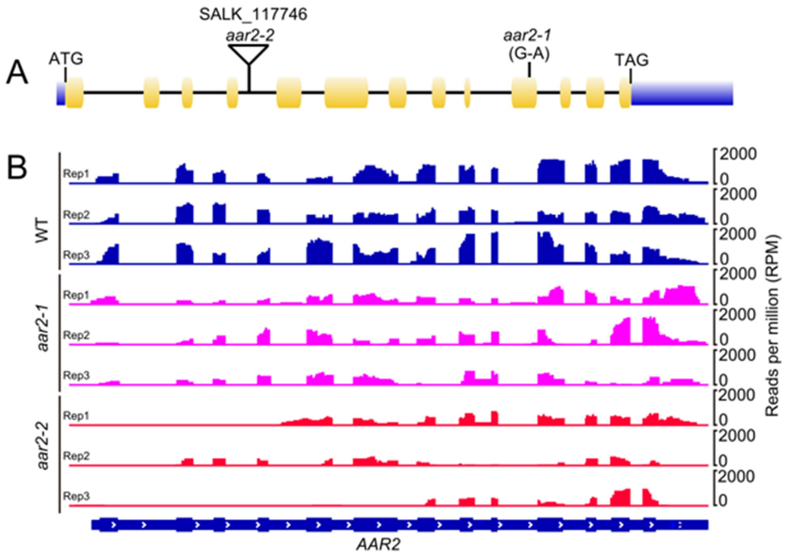


**Fig. S1. Identification of *aar2-1* and *aar2-2* mutations.** (*A*) A diagram of the *AAR2* gene showing mutation sites of *aar2-1* and *aar2-2.* Exons and introns are represented with rectangles and lines, respectively. The coding region and the UTRs are marked with yellow and blue rectangles, respectively. (*B*) *AAR2* expression in WT and *aar2-1* and *aar2-2* alleles as revealed by RNA-seq. Rep, replicates.

### Figure S2

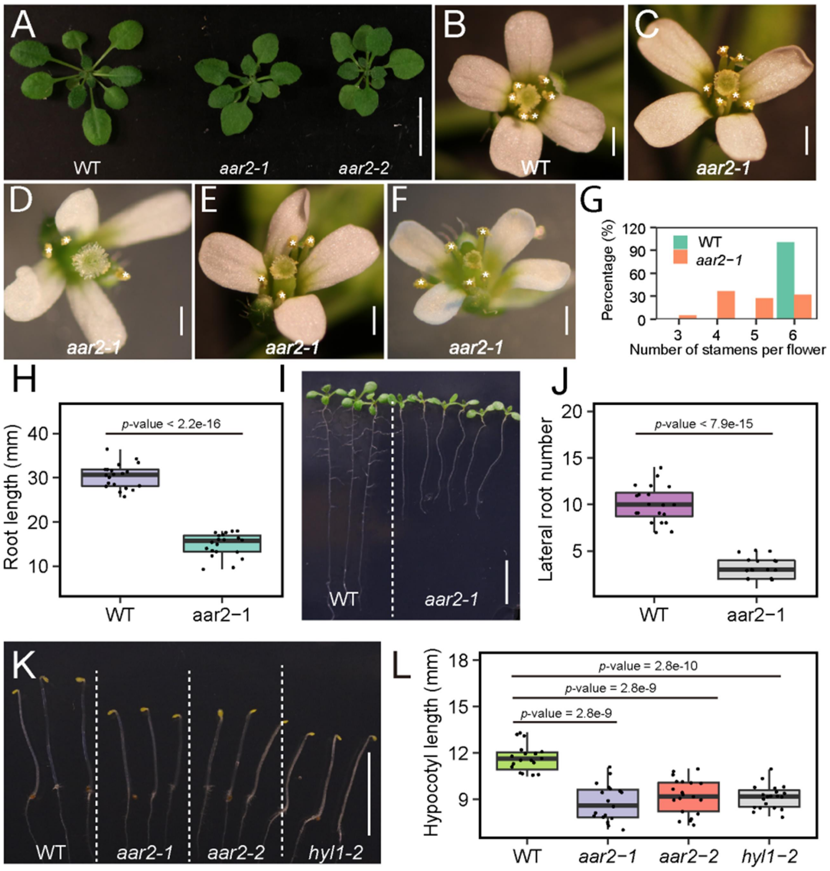


**Fig. S2. Developmental phenotypes of WT, *aar2-1* and *aar2-2* plants.** (*A*) 3-week old WT, *aar2-1* and *aar2-2* plants. (*B-F*) Top views of flowers in WT and *aar2-1* plants showing the reduced stamen number in some *aar2-1* flowers. (*G*) Quantification of stamen number in WT and *aar2-1* flowers. Percentages of flowers with 3, 4, 5, and 6 stamens per flower are shown. (*H*) Quantification of primary root length in WT and *aar2-1*. (*I*) 12-day-old seedlings of WT and *aar2-1* showing root length differences. (*J*) Quantification of lateral root number in WT and *aar2-1.* (*K*) Representative images of WT, *aar2-1*, *aar2-2*, *hyl1-2* grown in darkness. (*L*) Hypocotyl lengths of seedlings grown in the dark. *p*-values were calculated by student's t-test. Scale bar, 10 mm in (*A*), (*I*) and (*K*), 1 mm in (*B-F*).

### Figure S3

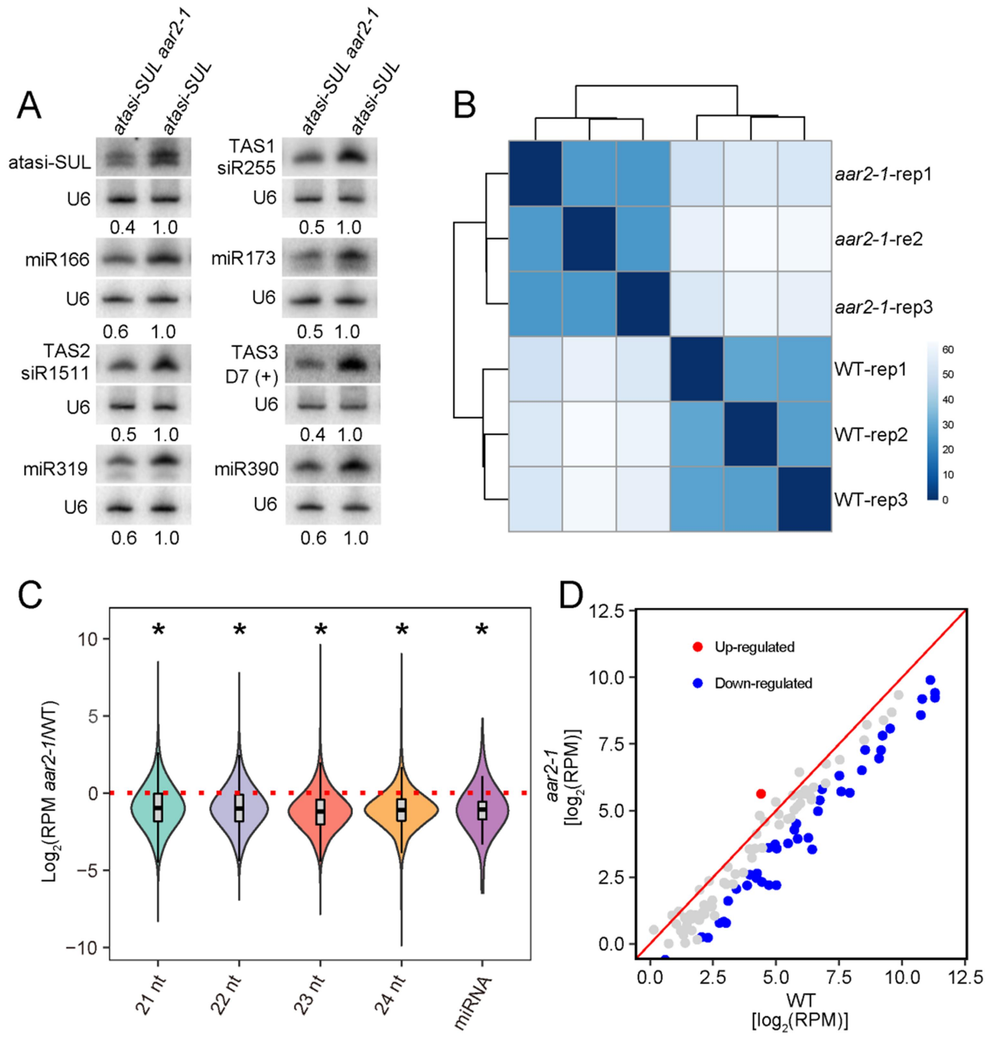


**Fig. S3. The *aar2-1* mutation leads to a global reduction in miRNA levels.** (*A*) RNA gel blot analysis showing reduced accumulation of atasi-SUL, endogenous ta-siRNAs and miRNAs in the *atasi-SUL aar2-1* mutant. The numbers represent relative levels. U6 was the loading control. (*B*) Clustering analysis showing that the three biological replicates are highly correlated. (*C*) Global abundance of 21-24 nucleotide (nt) small RNAs and all miRNAs in WT and *aar2-1* plants as determined by small RNA-seq. *P < 2.2e-16 as calculated by Wilcoxon test. (*D*) Scatter plots showing differentially accumulated miRNAs in WT and *aar2-1* mutants.

### Figure S4

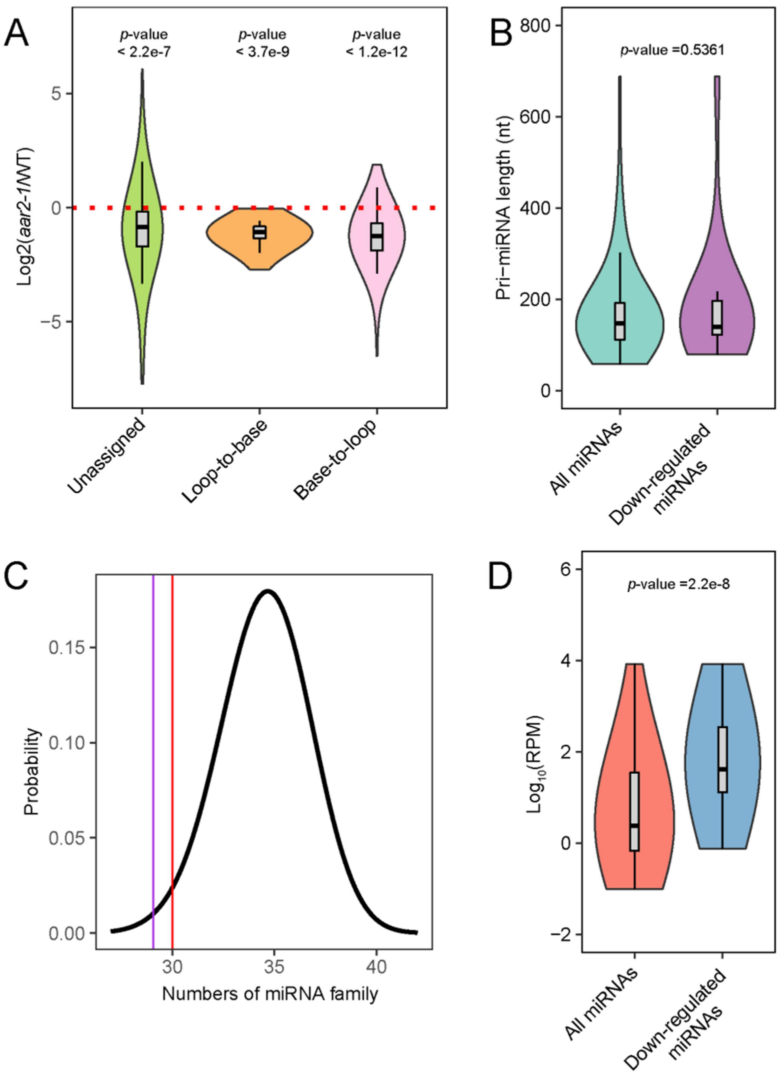


**Fig. S4. Common features of down-regulated miRNAs in *aar2-1*.** (*A*) Differential accumulation of miRNAs with different processing directions in *aar2-1* as compared with WT. All detected miRNAs were classified into three groups according to the direction of DCL1 processing of pri-miRNAs: loop-to-base, base-to-loop, and unassigned, according to published literature (1). (*B*) pri-miRNA length in all detected miRNAs and down-regulated miRNAs in *aar2-1.* (*C*) Comparison of the number of miRNA families represented by the 39 down-regulated miRNAs in *aar2-1* with that of randomly selected 39 miRNAs (n = 10000). The red line denotes the number of families represented by randomly selected miRNAs at a probability of 0.01. The blue line shows that the number of miRNA families represented by the 39 down-regulated miRNAs in *aar2-1* is 29. (*D*) Levels of all detected miRNAs and the 39 down-regulated miRNAs in *aar2-1*.

### Figure S5

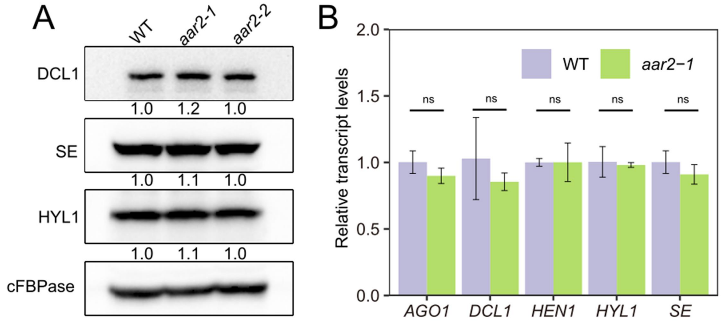


**Fig. S5. AAR2 does not influence the expression of key players in miRNA biogenesis.** (*A*) Protein levels of SE, DCL1, and HYL1 in WT, *aar2-1* and *aar2-2*. cFBPase was used as the loading control. Numbers represent protein levels relative to WT. (*B*) qRT-PCR analysis of the transcript levels of genes required for miRNA biogenesis or activities. Error bars represent SD calculated from three independent replicates. ns, not significant.

### Figure S6

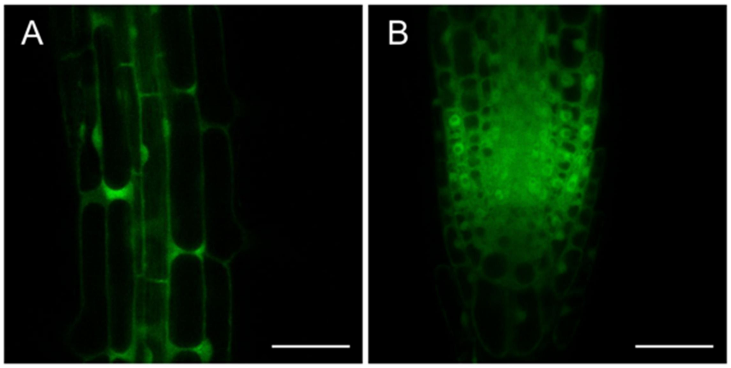


**Fig. S6. AAR2 is localized in both the cytoplasm and the nucleus.** (*A*) Subcellular localization of AAR2-YFP in the elongation zone of *AAR2::AAR2-YFP* transgenic roots. (*B*) Subcellular localization of AAR2-YFP in the root tip. Scale bar, 50 μm.

### Figure S7

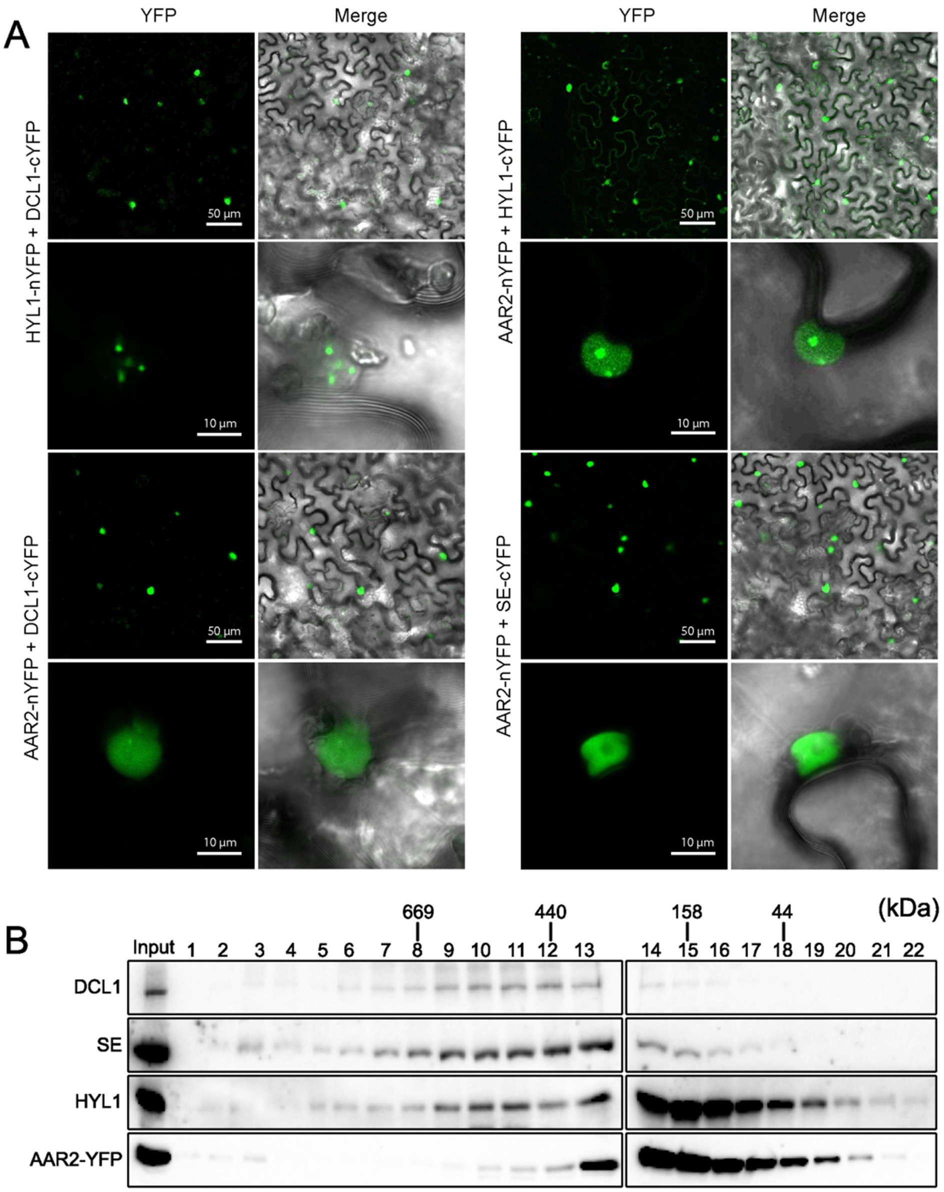


**Fig. S7. AAR2 associates with DCL1, SE, and HYL1.** (*A*) BiFC analysis showing the association between AAR2 and DCL1, SE, or HYL1. Paired cYFP- and nYFP-fusion constructs were co-infiltrated into tobacco leaves. The BiFC signals were detected at 48 h after infiltration by confocal microscopy. Scale bar, 50 μm in the first and third rows, 10 μm in the second and fourth rows. (*B*) Gel filtration analysis showing the co-distribution of AAR2 with DCL1, SE, and HYL1 in high molecular weight fractions (9-13). Protein lysates from *AAR2::ARR2-YFP* transgenic plants were fractionated according to size, followed by protein gel blot detection of DCL1, SE, HYL1 and AAR2-YFP. The fraction numbers are indicated above the gel images.

### Figure S8

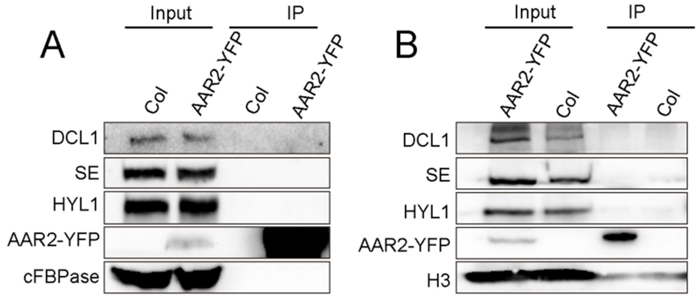


**Fig. S8. Co-immunoprecipitation (co-IP) to detect the interaction between AAR2-YFP and HYL1, SE and DCL1.** The western blots show the levels of various proteins in input and IP samples. cFBPase and histone H3 (H3) are loading controls. (*A*) Co-IP assay to examine the interaction between AAR2-YFP and HYL1, SE and DCL1 in total cell extracts. (*B*) Co-IP assay to examine the interaction between AAR2-YFP and HYL1, SE and DCL1 in nuclear extracts.

### Figure S9

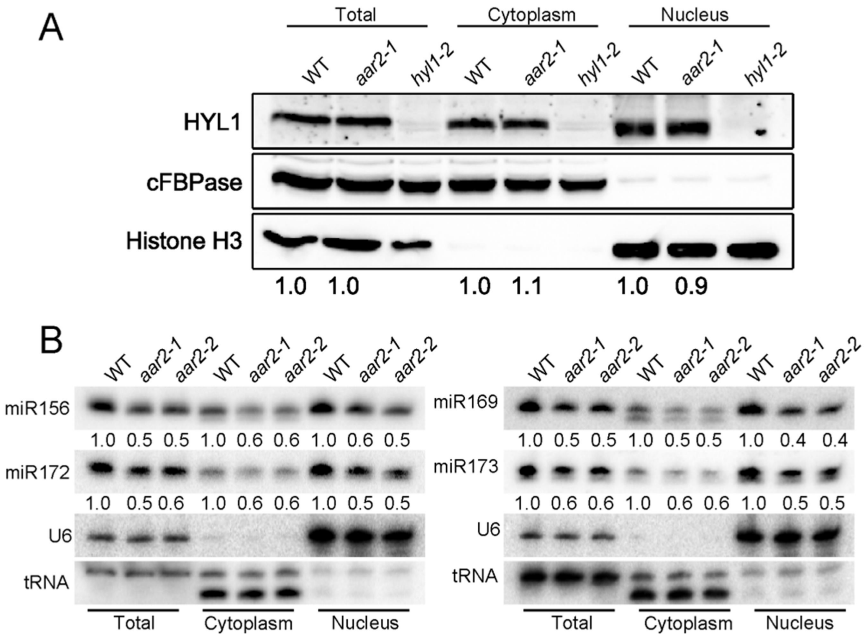


**Fig. S9. Subcellular distribution of HYL1 and miRNAs.** (*A*) Protein gel blot analysis showing the nucleo-cytoplasmic partitioning of HYL1. cFBPase was used as a cytoplasmic marker and histone H3 was used as a nuclear marker. HYL1 levels in *aar2-1* relative to WT are indicated by the numbers below the blots. (*B*) RNA gel blot analysis of miRNA accumulation in total, cytoplasmic and nuclear fractions in WT, *aar2-1* and *aar2-2.* U6 and tRNA were used as the nuclear and cytoplasmic markers, respectively, and as loading controls. miRNA levels were normalized against tRNA for the cytoplasmic fraction and U6 for the total and nuclear fractions.

### Figure S10

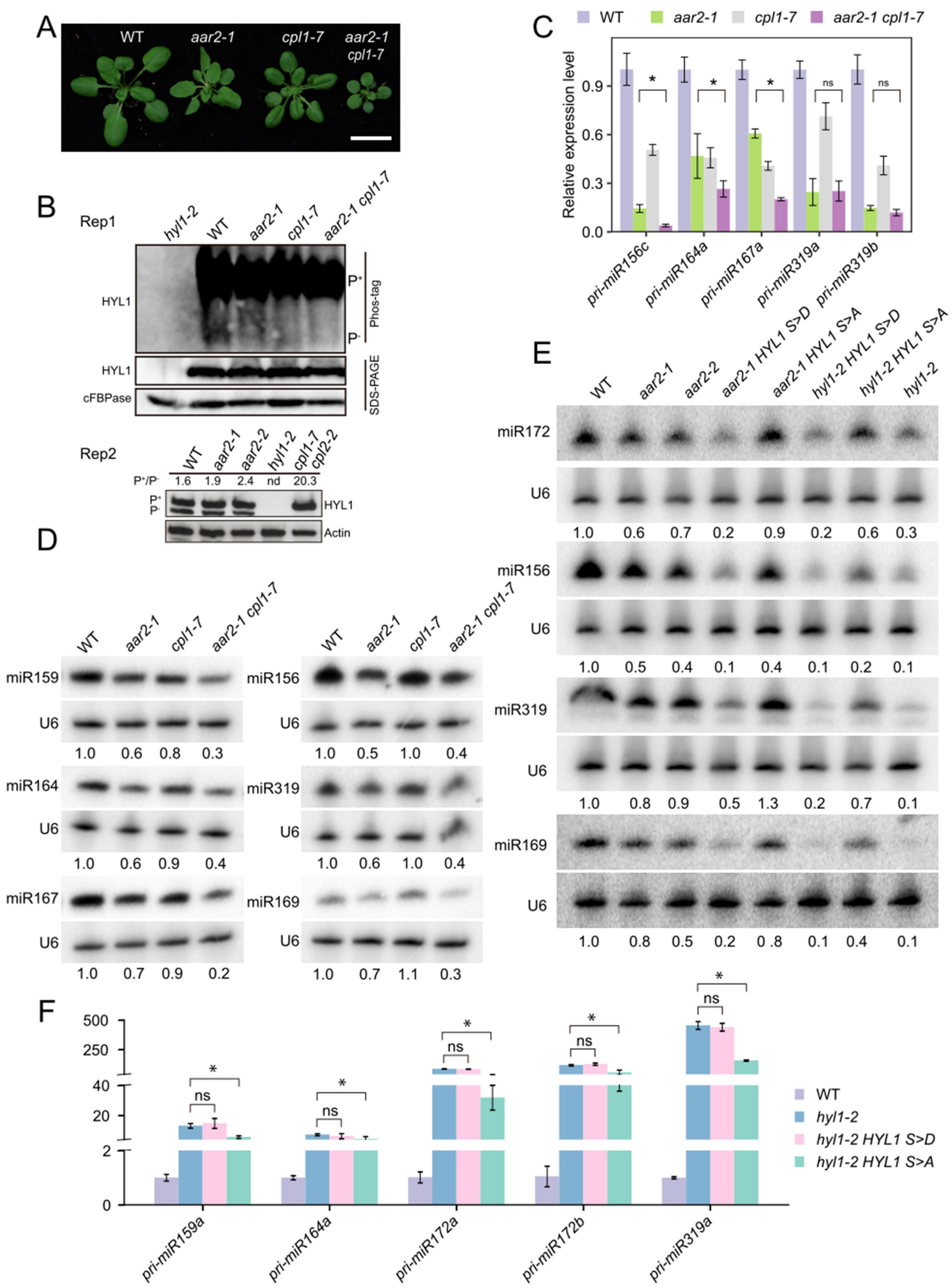


**Fig. S10. AAR2 impacts HYL1's phosphorylation status.** (*A*) 3-week-old plants of WT, *aar2-1, cpl1-7* and *aar2-1 cpl1-7*. (*B*) Phos-tag gel electrophoresis to detect phosphorylated ($P^+$) and non-phosphorylated ($P^-$) HYL1 in the indicated genotypes. cFBPase and actin were used as the loading controls. Rep1 (replicate 1), plants were grown in darkness for 5 days, transferred to continuous light and grown for another 3 days before analysis. Rep2, 14-day-old long-day grown plants were used for the analysis. Note that the $P^+/P^-$ ratios in wild type are different among Fig. 6A, Rep1, and Rep2. This is probably due to the different plant growth conditions: plants for Fig. 6A were grown under continuous light (70 μmol $m^{-2}$ $s^{-1}$) while those in Rep1 were grown under long-day (70 μmol $m^{-2}$ $s^{-1}$) before analysis; the plants in Rep2 were grown in Santa Fe, Argentina under long-day conditions using walk-in grown chambers with fluorescent lamps not enriched in red wavelength at an intensity of 150 μmol $m^{-2}$ $s^{-1}$. Fluctuations in the phosphorylation ratios between grown conditions/labs were previously observed (2-6) (*C*) qRT-PCR analysis of pri-miRNAs in WT, *aar2-1, cpl1-7* and *aar2-1 cpl1-7*. *UBQ5* served as an internal control. Error bars represent SD calculated from three independent replicates. Asterisks indicate significant difference (Student's t test, *P < 0.05). ns, no significant significance. (*D*) RNA gel blot analysis of miRNAs in WT, *aar2-1*, *cpl1-7* and *aar2-1 cpl1-7*. U6 was used as the loading control. The numbers represent relative miRNA levels. (*E*) RNA gel blot analysis of miRNAs in WT, *aar2-1, aar2-2, aar2-1 HYL1 S>D, aar2-1 HYL1 S>A, hyl1-2 HYL1 S>D, hyl1-2 HYL1 S>A,* and *hyl1-2*. U6 was used as the loading control. The numbers represent relative miRNA levels. (F) qRT-PCR analysis of pri-miRNAs in WT, *hyl1-2, hyl1-2 HYL1 S>D* and *hyl1-2 HYL1 S>A*. Asterisks indicate significant difference (Student's t test, *P < 0.01); ns, not significant.

### Figure S11

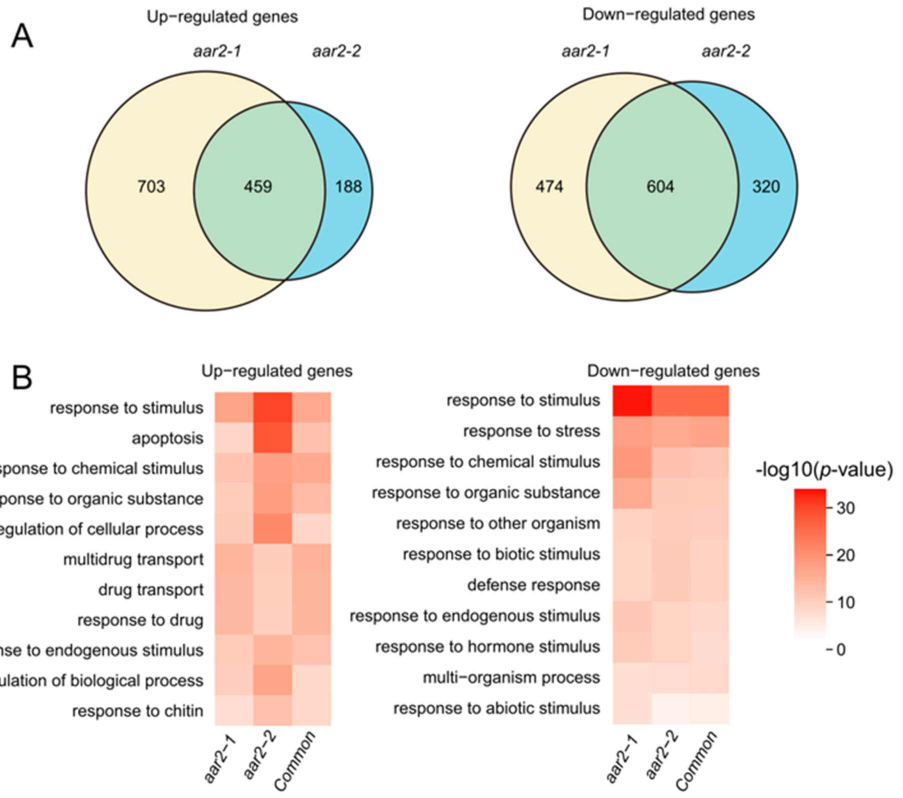


**Fig. S11. Differentially expressed genes in *aar2* mutants are enriched in those involved in responses to stimuli.** (*A*) Venn diagrams showing the overlaps in up-regulated genes and down-regulated genes between *aar2-1* and *aar2-2*. (*B*) Gene ontology enrichment analysis of up-regulated, down-regulated and commonly up-regulated and down-regulated genes between *aar2-1* and *aar2-2*.

### Figure S12

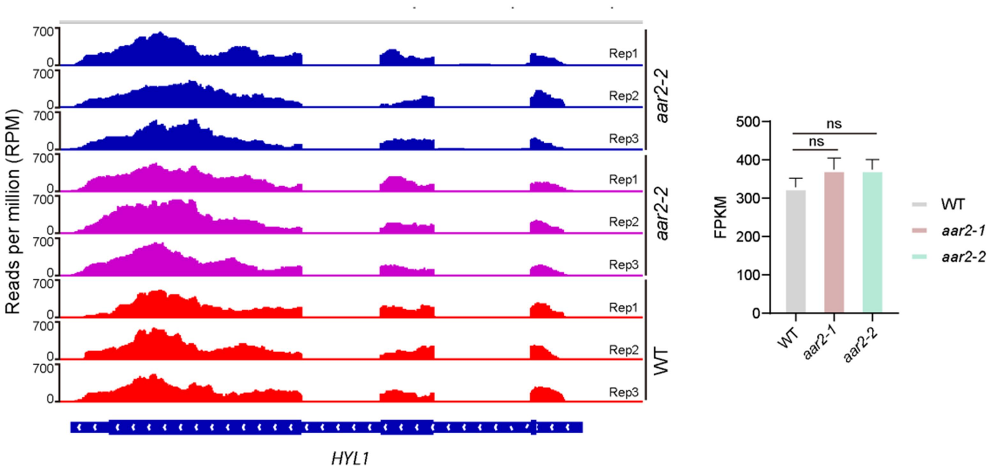


**Fig. S12.** (*A*) IGV tracks of *HYL1* in WT, *aar2-1* and *aar2-2* RNA-seq. rep, replicates. (*B*) Quantification of *HYL1* expression levels in WT, *aar2-1* and *aar2-2* by RNA-seq.

### Figure S13

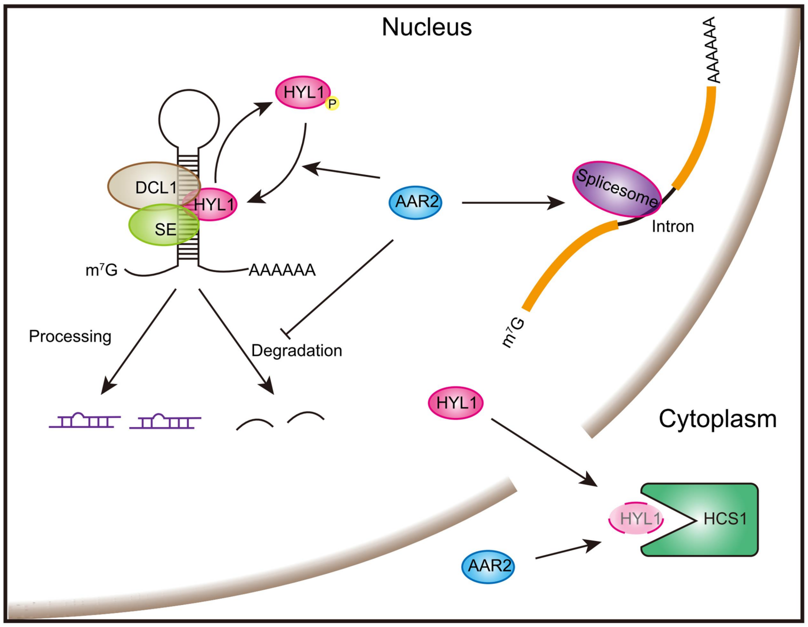


**Fig. S13. A proposed model for AAR2's functions in miRNA biogenesis and pre-mRNA splicing.** AAR2 promotes miRNA biogenesis by promoting HYL1 dephosphorylation and protecting pri-miRNAs from degradation. AAR2 also enables HYL1 degradation in the cytoplasm. AAR2 is present in both the nucleus and the cytoplasm.

complex in *Arabidopsis*. *Nat. Plants* 6(8):957-969.
